# Human genome-edited hematopoietic stem cells phenotypically correct Mucopolysaccharidosis type I

**DOI:** 10.1101/408757

**Authors:** Natalia Gomez-Ospina, Sam Glynne Scharenberg, Nathalie Mostrel, Rasmus O. Bak, Sruthi Mantri, Rolen M. Quadros, Channabasavaiah B. Gurumurthy, Ciaran Lee, Gang Bao, Laure Aurelian, Matthew H. Porteus

## Abstract

Lysosomal enzyme deficiencies comprise a large group of genetic disorders that generally lack effective treatments. A potential treatment approach is to engineer the patient’s own hematopoietic system to express high levels of the deficient enzyme, thereby correcting the biochemical defect and halting disease progression. Here, we present an efficient ex *vivo* genome editing approach using CRISPR/Cas9 that targets the lysosomal enzyme iduronidase to the *CCR5* safe harbor locus in human CD34+ hematopoietic stem and progenitor cells. The modified cells secrete supra-endogenous enzyme levels, maintain long-term repopulation and multi-lineage differentiation potential, and can correct biochemical and phenotypic abnormalities in an immunocompromised mouse model of Mucopolysaccharidosis type I. Our studies provide support for the development of human, genome-edited CD34+ hematopoietic stem and progenitor cells for the treatment of a multi-systemic lysosomal storage disorder. Our safe harbor approach constitutes a flexible platform for the expression of lysosomal enzymes, exemplifying a potential new paradigm for the treatment of these diseases.

## Introduction

Lysosomal storage diseases (LSDs) comprise a large group of genetic disorders caused by deficiencies in lysosomal proteins; many lack effective treatments. Mucopolysaccharidosis type I (MPSI) is a common LSD caused by insufficient iduronidase (IDUA) activity that results in glycosaminoglycan (GAG) accumulation and progressive multi-systemic deterioration that severely affects the neurological and musculoskeletal systems^1^. Current interventions for MPSI include enzyme replacement therapy (ERT) and allogeneic hematopoietic stem cell transplantation (allo-HSCT); both have limited efficacy. ERT does not cross the blood-brain barrier, requires costly life-long infusions, and inhibitory antibodies can further decrease enzyme bioavailability^2^. Allo-HSCT results in better outcomes than ERT by providing a persistent source of enzyme and tissue macrophages that can migrate into affected organs, including the brain, to deliver local enzyme^3-5^. However, allo-HSCT has significant limitations, including the uncertain availability of suitable donors, delay in treatment (allowing for irreversible progression), and transplant-associated morbidity and mortality such as graft-versus-host disease and drug-induced immunosuppression.

Human and animal studies in MPSI have shown that the therapeutic efficacy of HSCT can be enhanced by increasing the levels of circulating IDUA. In humans, patients transplanted with non-carrier donors had better clinical responses than patients transplanted with HSPCs from MPSI heterozygotes with decreased enzyme expression^6^. In mice, transplantation of virally transduced murine hematopoietic stem and progenitor cells (HSPCs) expressing supra-normal enzyme levels^7,8^ dramatically corrected the phenotype. Based on this, autologous transplantation of lentivirus-transduced HSPCs overexpressing lysosomal enzymes is being explored in human trials for LSDs^9^ (ClinicalTrials.gov, NCT03488394). This autologous approach eliminates the need to find immunologically matched donors and minimizes many of the potential complications from allogeneic transplants. However, concerns remain about the potential for tumorigenicity associated with random insertion of the viral genomes^10,11^, carry-over of infectious particles^12^, the immune response to some of the vectors, and variable transgene expression^13^.

Recently developed genome editing tools combine precise gene addition with genetic alterations that can add therapeutic benefit^14^. Among these, Clustered Regularly Interspaced Short Palindromic Repeats-associated protein-9 nuclease (CRISPR/Cas9) is the simplest to engineer and has been used to successfully modify HSPCs in culture^15^. The system was repurposed for editing eukaryotic cells by delivering the Cas9 nuclease, and a short guide RNA (sgRNA). When targeted to the sequence determined by the sgRNA, Cas9 creates a double-stranded DNA break, thereby stimulating homologous recombination with a designed donor DNA template that contains the desired genetic modification embedded between homology arms centered at the break site. This process, termed “homologous recombination-mediated genome editing” (HR-GE) is most often used for in-situ gene correction and has been hailed as a tool to treat monogenic diseases. Although its therapeutic potential in LSDs is unknown, to maximize therapeutic correction by autologous transplantation of genetically modified HSPCs in LSDs, functional enzymes must be expressed at higher-than-endogenous levels. This can be achieved by inserting an expression cassette (exogenous promoter-gene of interest) into non-essential genomic region (or “safe harbor”). A safe harbor provides a platform that is independent of specific patient mutations, is easily adaptable to various lysosomal enzymes and, compared to lentiviral transduction, ensures more predictable and consistent transgene expression because the insertion sites are restricted (up to 2 in autosomes).

Herein, we describe the development of such an approach for MPSI. We use *CCR5* as the target safe harbor to insert an expression cassette to overexpress IDUA in human CD34+ HPSCs and their progeny. *CCR5* is considered a non-essential gene because bi-allelic inactivation of *CCR5* (CCR5?32) has no general detrimental impact on human health and the only known phenotypes of CCR5 loss are resistance to HIV-1 infection and increased susceptibility to West Nile virus^16^. We report that human HSPCs modified using genome editing to express IDUA from the *CCR5* locus engraft and correct the biochemical, visceral, musculoskeletal, and neurologic manifestations of the disease in a new immunocompromised model of MSPI.

## Results

### Efficient targeting of IDUA into the *CCR5* locus in human HSPCs

To generate human CD34^+^ HPSCs overexpressing IDUA, we used sgRNA/Cas9 ribonucleoprotein (RNP) and adeno-associated viral vector serotype six (AAV6) delivery of the homologous templates^17^. RNP complexes consisting of 2′-*O*-methyl 3′phosphorothioate-modified *CCR5* sgRNA^18^ and Cas9 protein were electroporated into cord blood-derived (CB) and adult peripheral blood-derived HSPCs (PB). The efficiency of double-strand DNA break (DSB) generation by our *CCR5* RNP complex was estimated by measuring the frequency of insertions/deletions (Indel) at the predicted cut site. The mean Indel frequencies were 83% ± in CB-HSPCs and 76% ± 8 in PB-HSPCs, consistent with a highly active sgRNA. The predominant Indel was a single A/T insertion that abrogated CCR5 protein expression (**Extended Data Fig. 1**).

To achieve precise genetic modification, the templates for homologous recombination were made by inserting IDUA expression cassettes driven by the spleen focus-forming virus (SFFV) or the phosphoglycerate kinase (PGK) promoter, followed by a yellow fluorescent protein (YFP) downstream of the self-cleaving P2A peptide into the AAV vector genome. A third expression cassette containing IDUA driven by PGK but without a selection marker was also made (**Fig. 1a).** Following electroporation, CB and PB cells transduced with the SFFV-IDUA-YFP and PGK-IDUA-YFP viruses were examined for YFP fluorescence to quantify the efficiency of modification. As shown in **Figure 1b**, RNP electroporation followed by AAV6 transduction lead to a marked increase in the median fluorescence intensity of the cells. In CB-derived HSPCs the mean fraction of YFP-positive cells, was 34% ± 7 and 32% ± 8 with SFFV and PGK-driven expression cassettes respectively. In PB-HSPCs, the frequencies were 21% ± 5, and 24% ± 5 for the same AAV6 donors (**Fig. 1c).** AAV6 transduction alone showed <2% YFP positive cells, while mock cells that underwent electroporation but not AAV transduction had no detectable fluorescence. We measured the efficiency of modification in CB and PB cells transduced with the PGK-IDUA virus lacking the reporter (PGK-IDUA) by genotyping single cell-derived colonies from colony formation assays (CFAs) (**Extended Data Fig. 2a, b)**. In these cells, the frequencies of modification were 54% ± 10, and 44% ± 7 in CB and PB-HSPCs, considerably higher than the larger, YFP-containing cassettes, suggesting that efficiency is dependent on insert size (**Fig. 1c)**. Based on these targeting frequencies we conclude that our genome editing protocol is highly efficient and reproducible for human CB and PB-derived HSPCs.

**Fig. 1:**
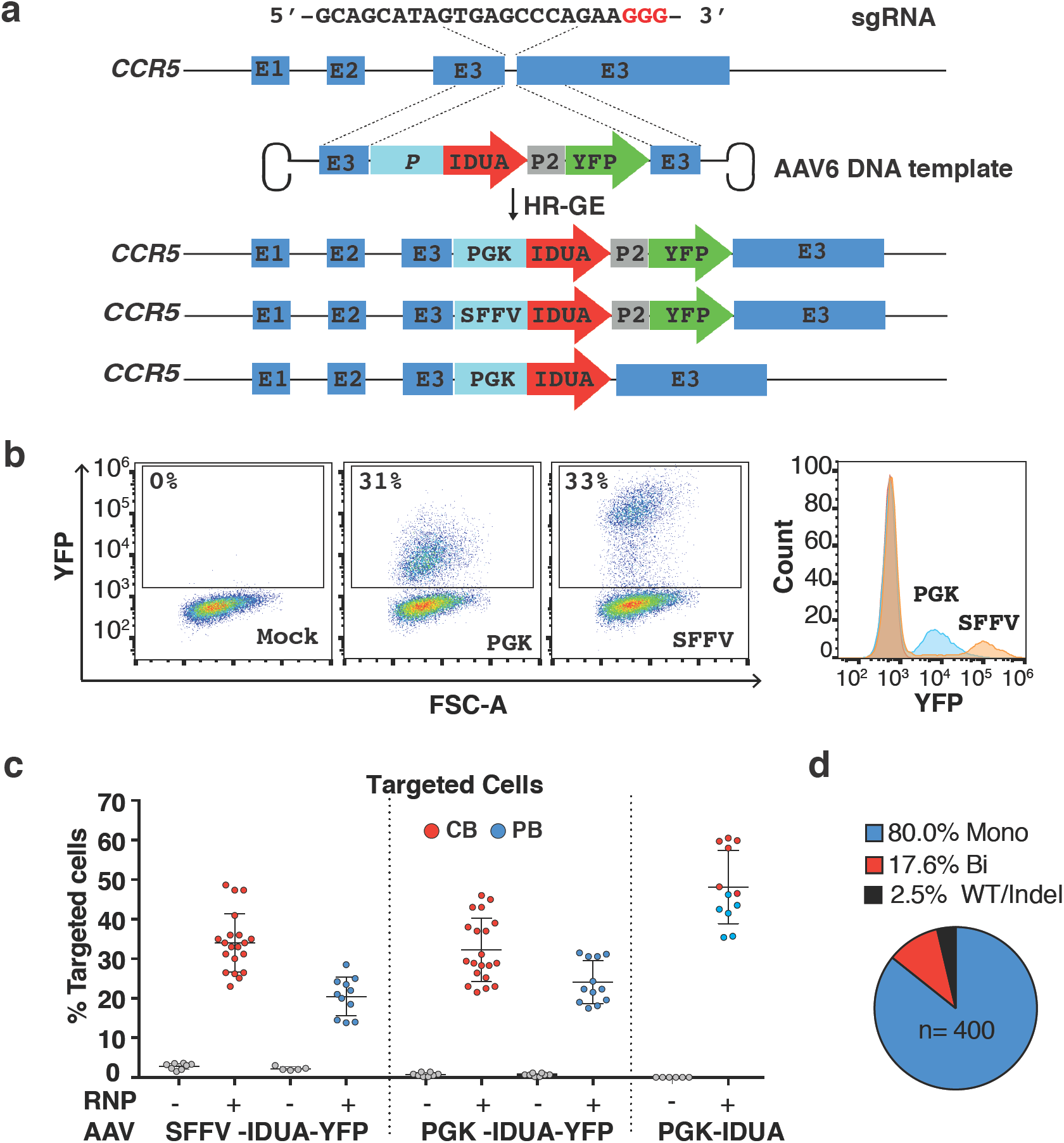
Efficient CRIPR/Cas9-mediated integration of IDUA overexpression cassettes into the *CCR5* locus in human CD34+ HSPCs. **a,** Schematic of targeted integration of IDUA and expression cassettes. The AAV6 genome was constructed to have 500bp arms of homology centered on the cut site, and the IDUA sequence placed under the control of the SFFV or the PGK promoter. In two DNA templates, YFP was expressed downstream of IDUA using the self-cleaving P2A peptide. Analysis was performed 3-days post-modification. **b,** FACs and histogram plots of mock and human HSPCs that underwent RNP and AAV6 exposure with YFP-containing expression cassettes. **c,** Targeting frequencies in CB (red) and PB (blue)-derived HSPCs read by percent fluorescent cells in YFP expressing cassettes and percent colonies with targeted CCR5 alleles by single cell-derived colony genotyping in cassettes without the reporter. Each dot represents the average of duplicates for a single human cell donor. For RNP+AAV6 conditions with YFP templates,CB=20, PB=11. For the template without selection CB=6, PB=6. Lines indicate mean and SD. **d,** Distribution of wildtype (WT), mono and bi-allelically modified cells in YFP-positive HSPCs (n=400, 3 human donors).

We also characterized the genomic modifications at the *CCR5* loci, by quantifying the fraction of targeted alleles in bulk DNA preparations using droplet digital PCR (ddPCR) **(Extended Data Fig. 2c, d)**. This data allowed us to estimate the distribution of cells with one (mono-allelic) or two (bi-allelic) alleles targeted and indicated that for the YFP constructs, 65% to 100% of the cells had mono-allelic modification **(Supplementary Data 1)**. Consistent with this, genotyping of YFP-positive colonies in CFAs showed an average mono-allelic modification frequency of 80% ± 7.5 (**Fig. 1d).**

### Enhanced IDUA secretion from edited HSPCs and HPSC-derived macrophages

A central concept in our approach is that HSPCs and their progeny will secrete stable, supra-endogenous IDUA levels that can cross-correct the lysosomal defect in affected cells. Examination of modified HSPCs in culture showed that 3 days post-modification, three distinct cell populations could be discerned based on YFP expression: high/medium/low (**Fig. 2a)**. YFP-high cells exhibited persistent fluorescence in culture for at least 30 days, demonstrating stable integration of the cassettes. YFP-negative cells had no detectable YFP expression at the time of selection, though 1% of cells eventually became positive. Most cells with intermediate fluorescence converted to YFP-high (80%) (**Fig. 2b)**. In these cultures with mixed YFP-positive and negative cells, grown under expansion conditions, the fraction of YFP-positive cells remained stable for 30 days, suggesting that neither the modification, nor the overexpression of the enzyme, nor the reporter impacted the cells’ proliferative potential.

**Fig. 2:**
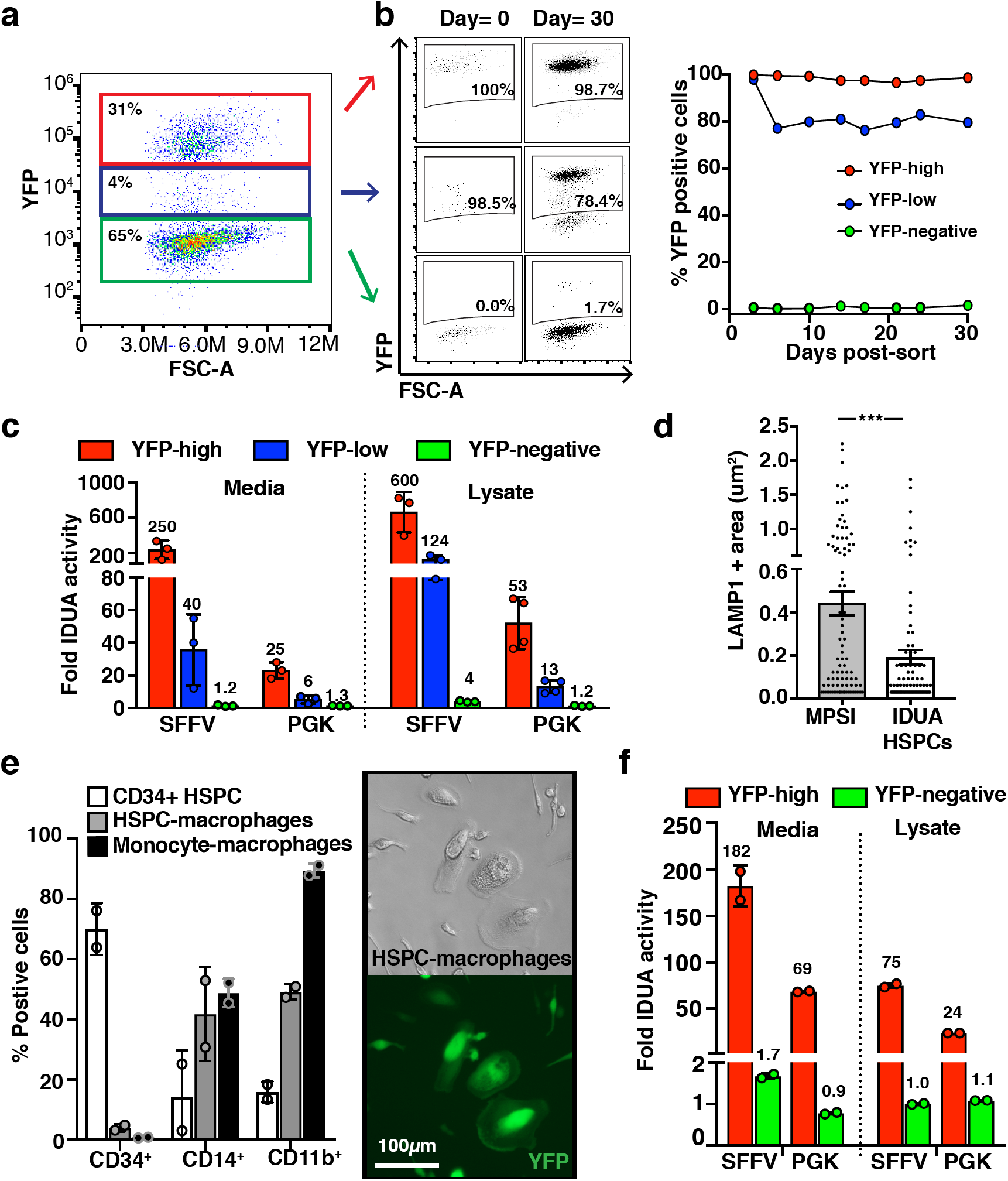
Enhanced IDUA expression by IDUA-HSPCs and derived macrophages. **a,** FACS plot shows distinct populations based on YFP expression 3 days post-modification. **b,** Persistent YFP expression up to 30 days cultures. **c,** Fold increase in IDUA secretion and intracellular expression by YFP-high, YFP-low, and YFP-negative populations compared to mock cells. **d,** Average LAMP-1+ area in MPSI fibroblasts co-cultured with IDUA-HSPCs. Each dot represents a cell. **e,** Human CD34, CD14, and CD11b marker expression in HSPC-derived macrophages and human monocyte-derived macrophages after *in vitro* differentiation compared to undifferentiated cells (CD34+ HSPCs). Macrophage morphology and YFP expression after differentiation. **f,** Fold increase in IDUA secretion and intracellular expression in HSPC-macrophages modified with SFFV and PGK expression cassettes. **c, e, and f,** Each dot represents average of triplicates in a human cell donor. All data expressed as mean ± SD, *** p < .001 in two-sided unpaired t-test.

When compared to mock-treated cells expressing endogenous IDUA levels, YFP-high cells secreted 250-fold and 25-fold more enzyme for the SFFV and PGK-driven cassettes respectively, while cell lysates expressed 600 and 50-fold more enzymatic activity (**Fig. 2c).** When YFP-high IDUA-HSPCs were co-cultured with patient-derived MPSI fibroblasts, they led to a decrease in the average area of lysosomal-associated membrane protein 1 (LAMP-1) positive specks, consistent with reduced lysosomal compartment size and cross-correction of the cellular phenotype (**Fig. 2d**). These data confirm that IDUA-HSPCs secrete supra-physiological IDUA levels and that the secreted IDUA has the post-translational modifications required for uptake into MPSI cells, thereby biochemically cross-correcting the patient derived enzyme deficient cells.

For IDUA-HSPCs to successfully correct biochemical abnormalities in the organs affected in MPSI, they must differentiate into monocytes that will migrate to and differentiate into tissue-resident macrophages such as microglia (brain), Kupffer cells (liver), osteoclasts (bone), and splenic macrophages to deliver the enzyme and cross-correct enzyme-deficient cells. To confirm that IDUA-HSPC could generate macrophages and that these cells can continue to produce IDUA, we differentiated these cells in culture and assayed for IDUA activity **(Supplementary Data 2 and Fig. 2e).** These IDUA-HPSC-derived macrophages secreted 182-fold and 69-fold more IDUA for the SFFV and PGK-driven cassettes respectively than mock-cell-derived macrophages. Likewise, lysates exhibited 75-fold and 24-fold more IDUA activity (**Fig. 2f).** These data established that IDUA-HPSC can reconstitute monocyte/macrophages *in vitro* and that IDUA-HPSC-derived macrophages also exhibit enhanced IDUA expression.

### Preserved repopulation and differentiation potential in IDUA-HSPCs

To determine if HSPCs that have undergone genome editing can engraft *in vivo*, we performed serial engraftment studies into NOD-scid-gamma (NSG) mice. We first tested cells modified with the SFFV and PGK constructs expressing YFP, which allowed us to identify the modified cells *in vivo*. Equal numbers of CB and PB-derived mock, YFP-negative (YFP-), and YFP-positive (YFP+) cells were transplanted intra-femorally into sub-lethally irradiated 6 to 8-week-old mice. Primary human engraftment was measured 16 weeks-post-transplantation by establishing the percent of bone marrow (BM) cells expressing both human CD45 and human leukocyte antigens (HLA-ABC) out of total mouse and human CD45+ cells (**Extended Data Figure 3b and Fig. 3a**). For the PGK-driven constructs, the median frequencies of hCD45^+^/HLA^+^ cells in BM were as follows: Mock 76.25% (min-max: 46.4-95.4%), YFP-21.5% (0.06-89.5%), YFP+ 4.3% (0.06-96%) (**Fig. 3b**). This showed a 5-fold drop in repopulation capacity in cells that underwent HR-GE (YFP+) compared to cells that did not but were also exposed to RNP, AAV transduction, and sorting (YFP-). The median frequency of human cells expressing YFP was 0.6% (0-18.5%) and 95.8% (1-100%) for YFP-and YFP+ transplants respectively, confirming that edited cells had engrafted in these mice (**Fig. 3c**). Human cells were also found in the peripheral blood with frequencies of 31/3.1/1.1% in mock, YFP-, and YFP+ cells respectively (**Fig. 3b)**.

**Fig. 3:**
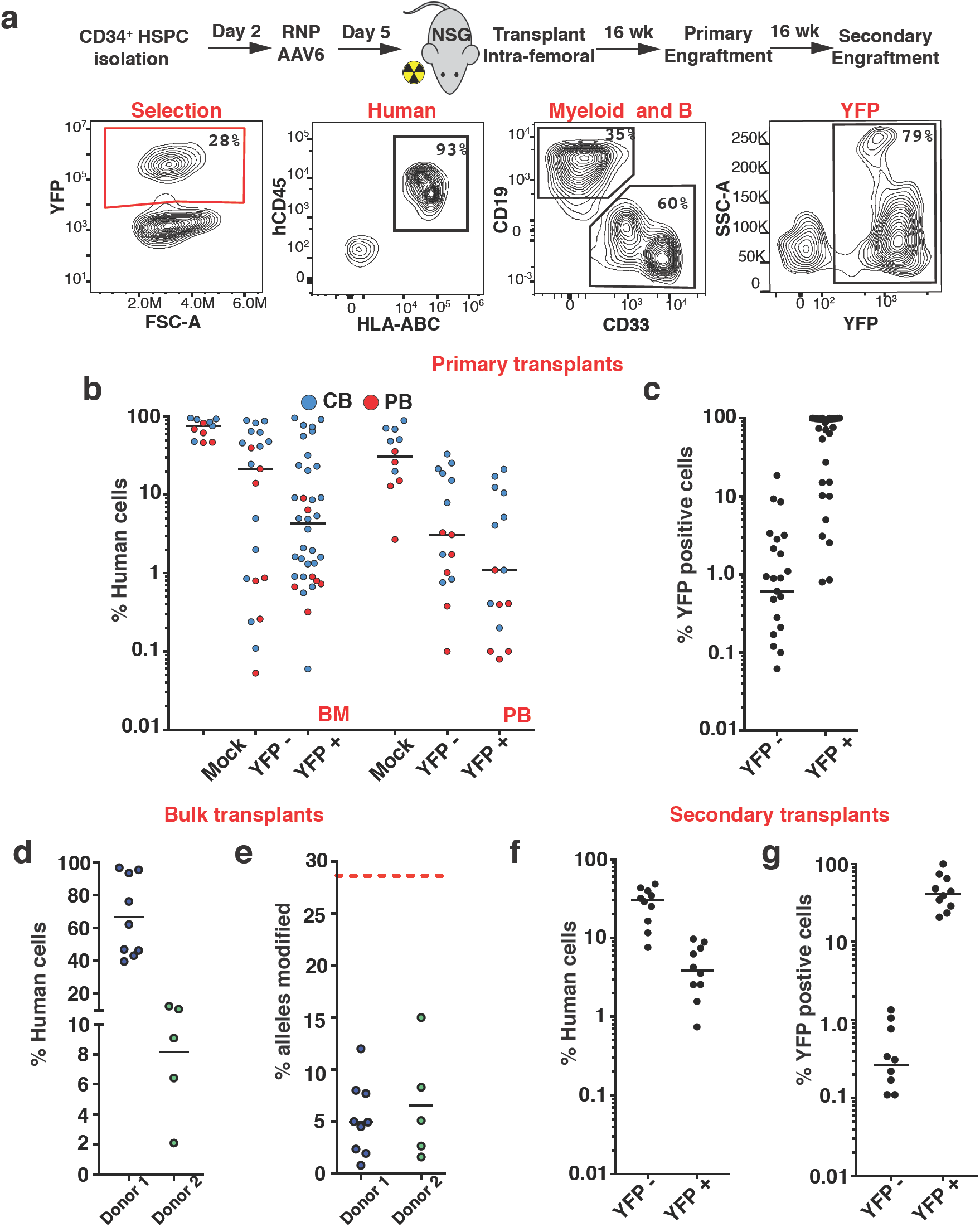
IDUA-HSPCs maintain long-term repopulation capacity. **a,** Schematic and representative FACS plots showing phenotyping by flow of human, myeloid, B-cell, and targeted cells after engraftment. **b,** Percent human cell chimerism in bone marrow (BM) and peripheral blood (PM) in mice 16-weeks post-transplant with CB (blue) and PB (red)-derived HSPCs targeted with PGK cassettes; mock (n=11), YFP-(n=21), and YFP+ (n=36). Each point represents a mouse. **c,** Percent human YFP+ cells in BM of mice in BM 16-weeks post-transplant. **d,**Percent human cell chimerism in BM in mice transplanted with bulk cells without selection from two different human cell donors. **e,** Percent modified alleles in engrafted cells by ddPCR. 28% was the starting allele modification frequency. **f,** Percent human cell chimerism in BM of mice in secondary transplants 32 weeks after genome editing; YFP-(n=10), and YFP+ (n=10). **g,** Percent human, YFP+ cells in BM of mice in secondary transplants.

The apparent engraftment advantage of cells that had not undergone HR-GE was also examined by transplanting bulk populations of HSPCs modified with the cassette without YFP. In two independent experiments, an initial fraction of targeted alleles of 28% (43% modified cells) declined to 5.2% and 6.5% in the engrafted cells (8 and 10% modified cells) despite big differences in human chimerism **(Fig. 3c, d).** This corresponded to a 5-fold drop in donor 1 and 4-fold drop in donor 2. Interestingly, this fall in targeted alleles showed significant variation in individual mice (2 to 10-fold). This data re-demonstrated the observed loss in engraftment potency after modification and supports the idea that the HR-GE cell population has fewer clones with long-term repopulation potential.

Serial transplantation is considered a gold standard to assess self-renewal capacity of HSCs. For secondary transplants, we isolated human CD34^+^ cells from the bone marrow of primary mice and transplanted into secondary mice. YFP+ engrafted mice showed 3.9% (0.8-9.7%) median human cell chimerism, while YFP-mice showed 30.4% (7.7-48.2%) (**Fig. 3e**). YFP expression in the engrafted human cells was 0.27% (0-1.35) for YFP-cells, and 41.9% (20.8-100) for YFP+ cells (**Fig. 3f)**. Similar levels of human cell chimerism were observed for the SFFV-driven constructs in serial transplants (**Extended data Fig. 4**). Collectively, the presence of YFP expressing cells at 16-and 32-weeks post-modification demonstrates that cells with long-term repopulation potential can be edited, albeit at lower frequencies than cells that did not undergo HR-GE.

To establish the modified cells’ ability to differentiate into multiple hematopoietic lineages, we looked *in vitro* using colony formation assays (CFAs) and *in vivo* after engraftment in NSG mice. In CFAs, CB-derived and PB-derived YFP-expressing cells gave rise to all progenitor cells at the same frequencies as mock treated and YFP-cells, indicating that IDUA-HSPCs can proliferate and differentiate into multiple lineage progenitors in response to appropriate growth factors (**Extended data Fig. 5a, b**). *In vivo*, B, T and myeloid cells were identified using the human CD19, CD3, and CD33 markers. Compared to mock cells that demonstrated a roughly equal distribution of B and myeloid cells (1:1, CD19:CD33) 16 post-transplantation, YFP+ and YFP-cells showed skewing towards myeloid differentiation (YFP+ = 1:16, and YFP-= 1:5) (**Extended data Fig. 5c)**. Subgrouping the mice based on human-cell chimerism showed that in mice with low human engraftment (< 1%), the cells exhibited myeloid skewing (72% of these mice had human myeloid cell fractions >90%). In contrast, mice with chimerism greater than 20% displayed a mean CD33+ fraction of 55% (**Extended Data Fig. 5d, e)**. This myeloid bias was not observed in circulating cells in the peripheral blood or in secondary transplants (**Extended data Fig. 5f, g**). These data suggest that myeloid skewing is tightly and inversely correlated with the degree of human cell engraftment, and that neither the genome editing process, nor IDUA expression, affects the modified cell’s capacity to differentiate into multiple hematopoietic lineages *in vitro* or *in vivo*.

### Biochemical correction in NSG-IDUA^X/X^ mice by human IDUA-HSPCs

To determine the potential of human IDUA-HSPCs to correct the metabolic abnormalities of MPSI, we established a new mouse model of the disease capable of engrafting human cells. NSG-IDUA^X/X^ mice replicated the phenotype of patients affected with MPSI^1^ and previously described immunocompetent^19,20^ and immunocompromised^21^ MPSI mice (**Supplementary Data 3, Extended data Fig. 6 and 7)**. We focused the correction experiments on cells expressing IDUA under the PGK promoter, as this promoter has better translational potential because it has decreased enhancer-like activity compared to SFFV^22^. As a first series of experiments, we examined PB-derived cells in which the modification did not include a selection marker. In bulk transplants, the median human cell chimerism in the bone marrow was 62.2% (min=39.2, max=96.7%) and no statistically significant differences in human engraftment were observed between NSG-IDUA^X/X^ and NSG-IDUA^W/X^ mice (**Fig. 4a**). The editing frequencies before transplantation were 30% of *CCR5* alleles (by ddPCR) and 46% of cells (as measured by CFA). GAG urinary excretion was measured at 4, 8, and 18 weeks post-transplantation in NSG-IDUA^X/X^ and IDUA^W/X^ mice. Biochemical correction was detectable after 4 weeks, and the trend towards normalization increased over time (**Fig. 4b**). These kinetics are consistent with the time lag needed for the genetically engineered human HSPCs to engraft, expand, and migrate to affected tissues and “cross correct” diseased cells. At 18 weeks, NSG-IDUA^X/X^ mice that had been transplanted with IDUA-HSPCs (X/X Tx) excreted 65% less GAGs in the urine compared to sham-treated NSG-IDUA^X/X^ mice (X/X sham) (median Tx= 387.2 µg/mg of creatinine, sham=1,122 µg/mg) though the levels had not normalized (W/X sham =155 µg/mg) (**Fig. 4b**). Transplantation of IDUA-HSPCs also resulted in increased IDUA activity to 11.3%, 50.1%, 167.5%, and 6.8% of normal in serum, liver, spleen, and brain respectively (compared to undetectable in X/X sham) and resulted in normalization of tissue GAGs in liver and spleen (**Fig. 4c, d**). Supra-endogenous levels of activity were detected consistently in the spleen and occasionally in the liver of some mice. This can be attributed to robust human cell engraftment in these organs, as demonstrated by increases in liver and spleen size in transplanted mice regardless of genotype (**Extended Data Fig. 8b-e)**.

**Fig. 4:**
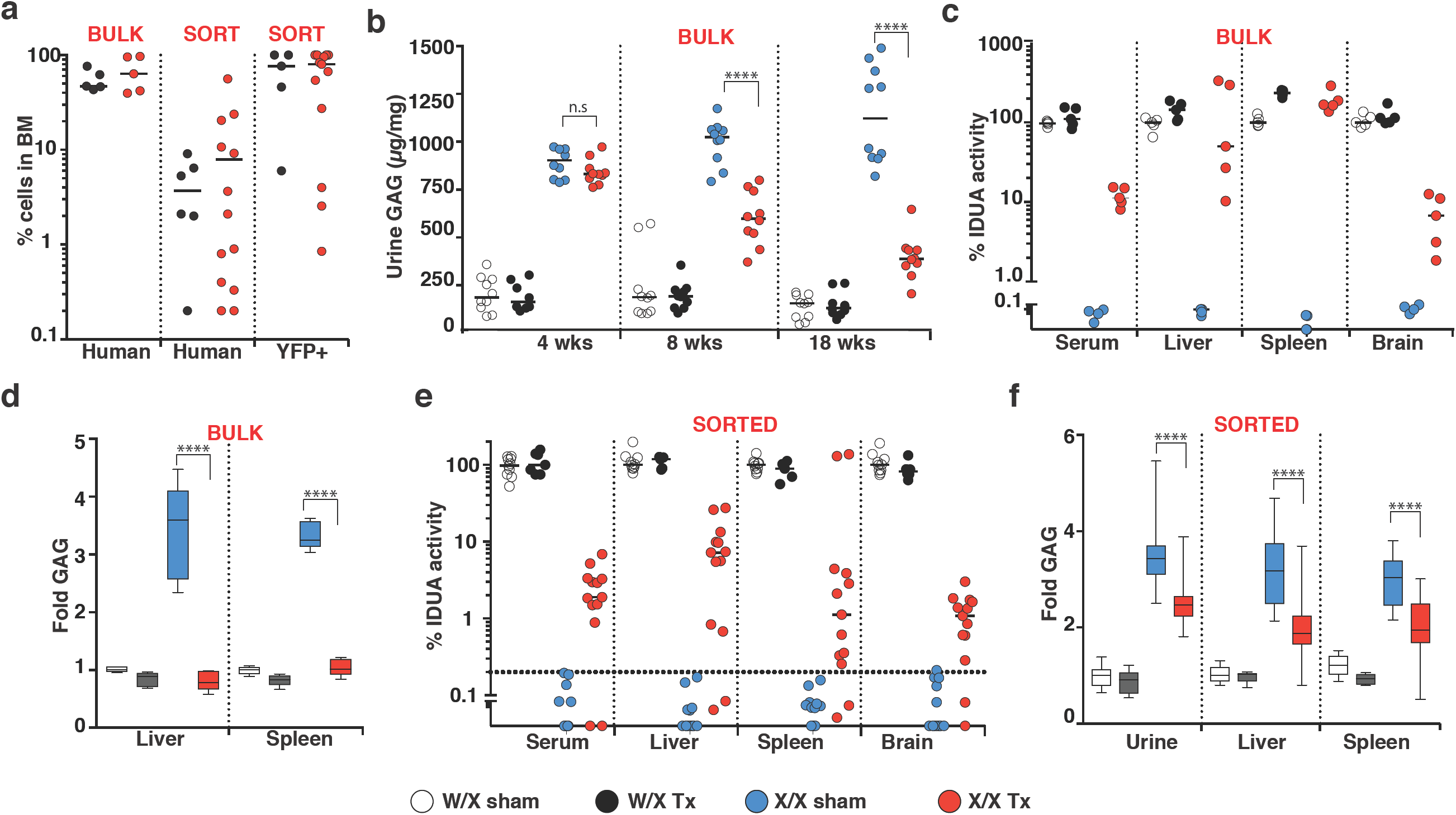
Biochemical correction in NSG-IDUAX/X mice by human IDUA-HSPCs. IDUA activity and GAG accumulation in heterozygous sham-treated (W/X sham-clear), heterozygous transplanted (W/X Tx-black), homozygous sham-treated (X/X sham-blue), and homozygous transplanted (X/X Tx-red) mice. **a,** Percent human and YFP+ cells in BM in experiments using bulk and sorted cells. **b,** Urinary GAGs at 4,8, and 18 weeks in experiments using bulk cells (n=5 mice per cohort, two measurements per mouse). **c,** Serum and tissue IDUA activity in experiments using bulk cells (n=5 per cohort). **d,** Fold GAG storage in liver and spleen (normalized by W/X sham, n=5 per cohort). **e,** Serum and tissue IDUA activity in experiments using sorted cells (n=5 for W/X Tx and sham mice, and n=13 for X/X Tx and sham mice). **f,** Fold GAG urinary excretion and tissue storage in experiments using sorted cells (normalized by W/X sham). Median values in all scatter plots. d and f show box plots with whiskers at the 5-95th percentiles. ****: p < .0001 in one-way ANOVA test. Post hoc comparisons were made with the Tukey’s multiple comparisons test.

Because we could not discount the contribution of unmodified cells to the observed correction in bulk transplants, we then examined the effect of HSPCs expressing IDUA and YFP under the PGK promoter after FACS. Of 15 NSG-IDUA^X/X^ and 5 NSG-IDUA^W/X^ 13/15 and 5/5 were deemed to have engrafted (human chimerism in the bone marrow >0.1%). The median percent human chimerism was 4.2% (W/X) and 9.9% (X/X). IDUA-YFP-HSCPs increased IDUA tissue activity to 2.9%, 7.4%, 25.5%, and 1.3% of normal in serum, liver, spleen, and brain respectively (**Fig. 4a)**. Tissue and urine GAGs were also significantly reduced (**Fig. 4f)**. Together, this data indicates that IDUA-HSPCs can correct the metabolic abnormalities in MPSI and suggest that the degree of correction correlates with human cell chimerism.

### Phenotypic correction in NSG-IDUA^IDUAX/X^ mice by human IDUA-HSPCs

To investigate the effect of IDUA-HSPCs on the skeletal and neurological manifestations of MPSI, sham-treated and transplanted mice also underwent whole body micro-CT and neurobehavioral studies 18 weeks after transplantation. The effect of transplantation on the skeletal system was measured on the skull parietal and zygomatic bone thickness and the cortical thickness and length of femoral bones. In experiments where the mice were transplanted using unselected cells and where human cell chimerism was high (**Fig. 4d**), we observed almost complete normalization of bone parameters by visual inspection and on CT scan measurements (**Fig. 5a, b**). Mice transplanted with cells that had undergone selection showed partial but statistically significant reduction in the thickness of the zygomatic, parietal bones, and femur (**Fig. 5c**).

**Fig. 5:**
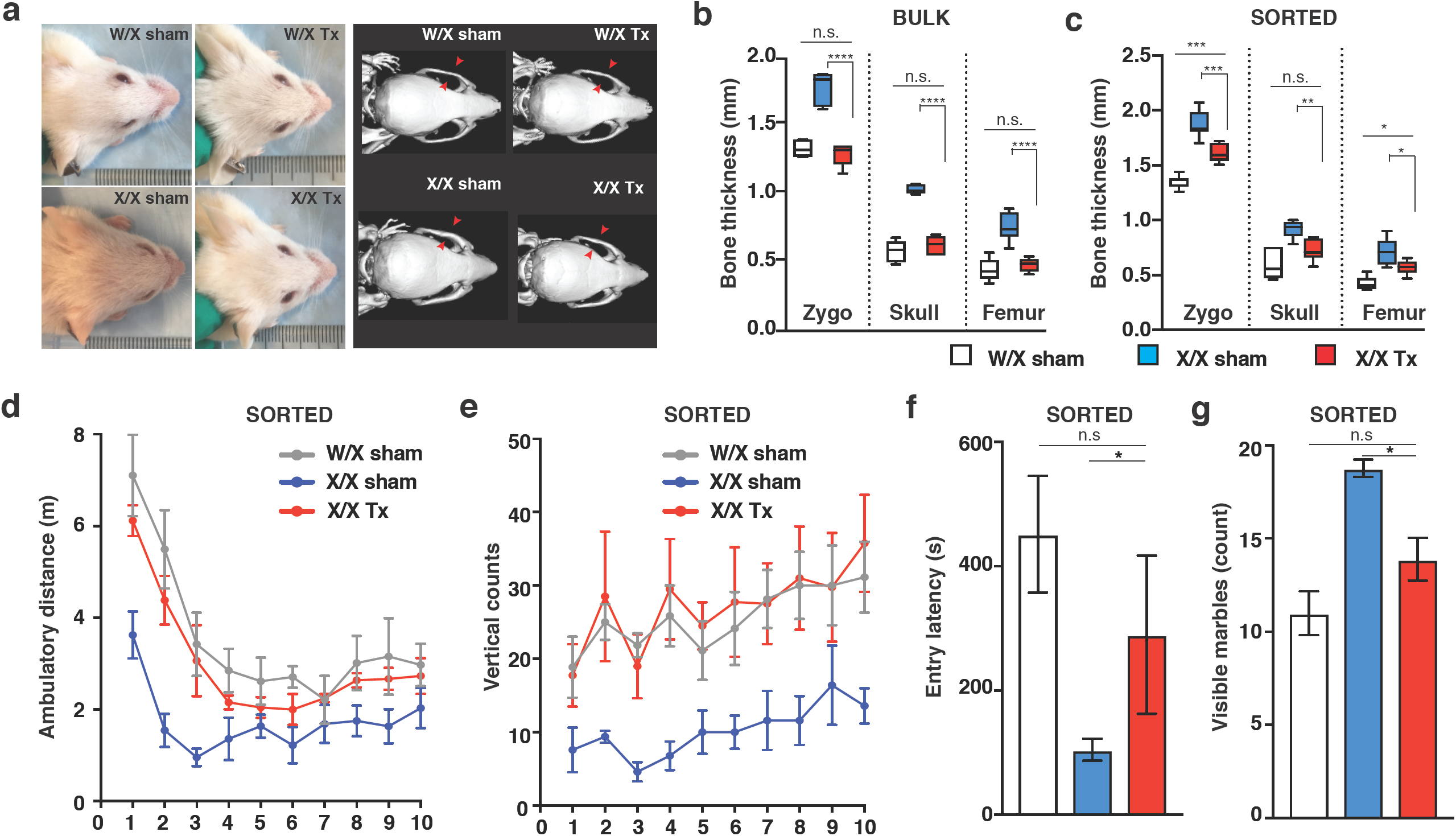
Phenotypic restitution in NSG-IDUAX/X mice by human IDUA-HSPCs. Behavioral and skeletal assessment in: W/X sham (clear or gray, n=11), X/X sham (blue, n=10), and X/X Tx (red, n=11). **a,** Representative photos showing facial features in mice transplanted with bulk cells. **b,** Bony features in mice transplanted with bulk and **c,** sorted cells. Box plots with whiskers show min and max. **d,** Ambulatory distance in mice transplanted with sorted cells. W/X sham vs. X/X sham: **; W/X sham vs. X/X Tx: n.s.; X/X sham vs. X/X sham: *. **e,** Vertical rearing in mice transplanted with sorted cells. W/X sham vs. X/X sham: *; W/X sham vs. X/X Tx: n.s.; X/X sham vs. X/X sham: *. **f,** Memory retention in mice transplanted with sorted cells. **g,** Quantification of digging behavior in mice transplanted with sorted cells. Data shown as mean ± SEM. **b-g,** Comparisons between groups were performed using one-way ANOVA test and post-hoc comparisons were made with the Tukey’s multiple comparisons test. *: p < .05, **: p < .01, ***: p < .001, and ****: p < .0001. Open field testing and vertical rearings were analyzed using within-subject modeling by calculating the are under the curve for each mouse within the first five minutes and comparing between groups with one-way ANOVA.

We also examined the open field behavior, passive inhibitory avoidance, and marble-burying behavior of sham-treated and transplanted mice. Transplantation of bulk-edited cells resulted in marked reduction in locomotor activity and long-term memory, regardless of genotype **(Extended Data Fig. 9a-c**). We suspected that high human-cell chimerism was detrimental for the overall health of the mice. Consistent with this, we observed growth restriction following human cell transplantation in both homozygous and heterozygous mice (**Extended Data Fig. 8a)**. This likely represents a toxicity artifact of this xenogeneic transplant model and could be explained in part by the defective erythropoiesis seen in these xenograft models^23^ when human cell chimerism is high. In contrast, NSG-IDUA^X/X^ mice transplanted with YFP-selected cells in which human cell chimerism was low exhibited locomotor activity indistinguishable from their sham-treated heterozygous littermates, and markedly higher that the sham-treated knock-out mice (**Fig. 5d**). These mice also had increased vertical counts at all time points and demonstrated the same exploratory behavior as sham heterozygous mice (**Fig. 5e**). Transplantation of IDUA-HSPCS in NSG-IDUA^X/X^ also enhanced performance in the passive inhibitory avoidance test 24 hours later (**Fig. 5f)**. Digging and marble burying behavior also improved but did not completely normalize (**Fig. 5g)**.

### Safety of our genome editing strategy

To assess genotoxicity and characterize the off-target repertoire of our *CCR5* guide, we used the bioinformatics-based tool COSMID (CRISPR Off-target Sites with Mismatches, Insertions, and Deletions)^24^. Off-target activity at a total of 67 predicted loci was measured by deep sequencing in two biological replicates of CB-derived HSPCs. In each replicate we compared the percent Indels measured in mock and cells electroporated with RNP with either wild-type (WT) Cas9 or a higher fidelity (HiFi) Cas9^25^. Five of the 67 sites were located within repetitive elements and Indel rates could not be assigned to specific loci in this group **(Extended Data Fig. 10)**. For the remaining 62 genomic locations, sites were deemed true off-targets if: 1) the percent of indels at the site was > 0.1% (limit of detection), 2) off-target activity was present in both biological samples, and 3) indels were higher in the RNP compared with the mock samples. Given these criteria only 4 sites were deemed to be true off-targets (**Figure 6a, b)**. For all of these sites the frequency of Indels was < 0.5% and the use of the HiFi Cas9 abolished off-target activity entirely while maintaining on-target efficiency. Only one exonic site was found in the *SUOX* gene (sulfite oxidase). The highest off-target activity measured at this site was 0.128%, which was reduced below the limit of detection with HiFi Cas9. These data suggest that our *CCR5* sgRNA combined with either WT Cas9 or especially HiFi Cas9 has negligible off-target activity on a large screen of bioinformatically predicted sites.

**Fig. 6:**
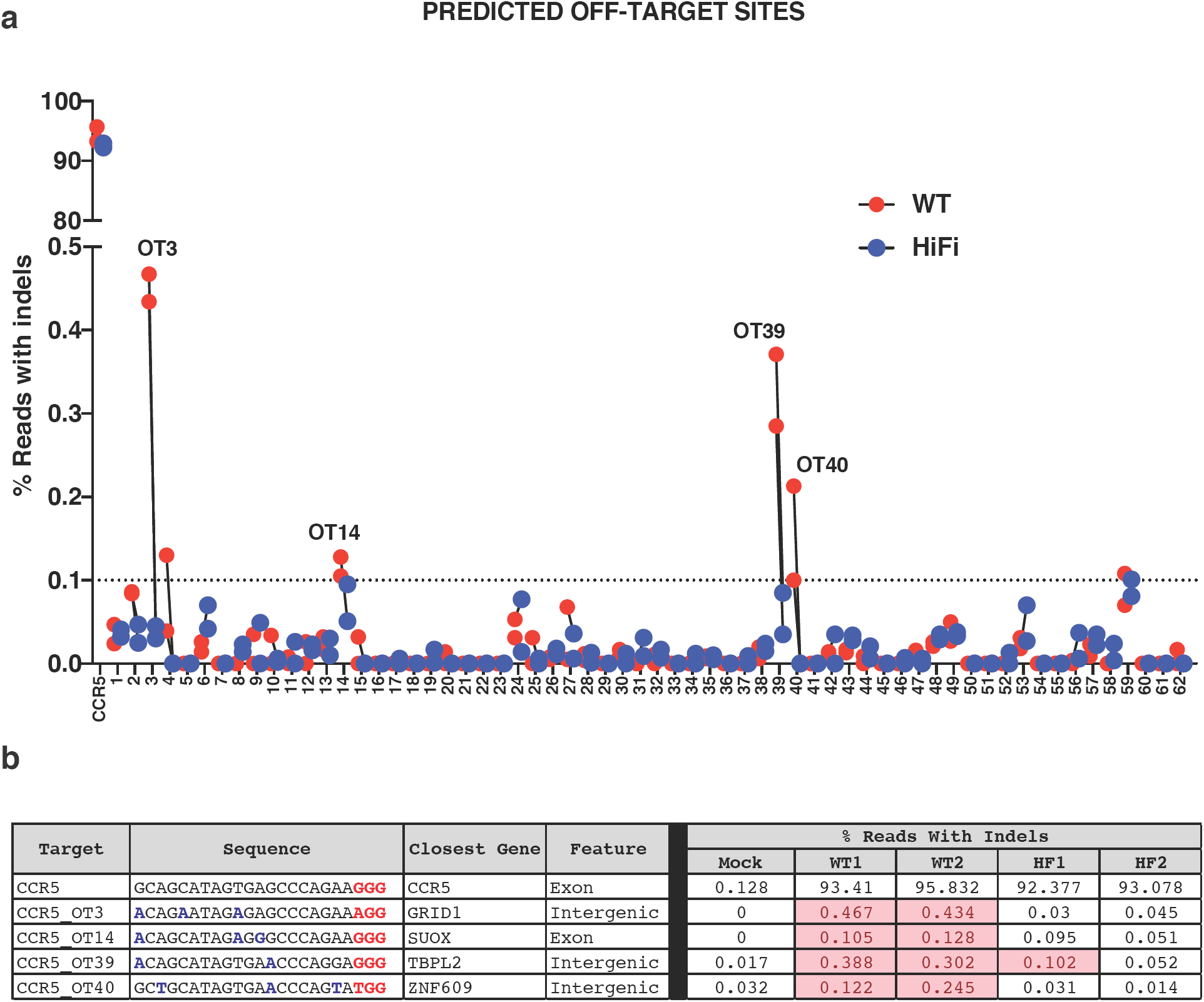
OFF-target analysis of the CCR5 sgRNA. **a,** Percent reads with Indels at 62 off-target sites predicted using COSMID. For each site, red dots indicate samples treated with WT Cas9 and blue dots indicate samples treated with HiFi Cas9. The limit of detection for NGS is 0.1% and is indicated on the graph by a dashed line. **b,** Table summarizing four bona fide off-target sites. PAM sequences are shown in red and mismatched bases are shown in blue. For all of these sites the percent of Indels was < 0.5%. For all of these sites, the use of the HiFi Cas9 abolished off-target activity.

Collectively, we performed 191 autopsies (101 mice used in primary engraftment, 50 in secondary engraftment, and 40 in NSG-IDUA^X/X^ correction studies) in which no gross tumors were found. Three tumor-like masses were evaluated by histology and confirmed to be abscesses. These 191 mice were transplanted with a combined dose of 90 million human cells that underwent our genome editing protocol. Considering that the median age for HSCT in MSPI patients is around one year^26^, and that an average one year-old is 10 Kg, the total number of modified cells used in this study is roughly equivalent to two clinical doses of 4.5 x 10^6^ CD34 HSPCs/kg. We conclude that the apparent lack of tumorigenicity and the low off-target activity of the *CCR5* sgRNA provide evidence for the safety our modification strategy.

## Discussion

We describe an efficient application of RNP and AAV6-mediated template delivery to overexpress IDUA from a safe harbor locus in human CD34+ HSPCs. The suitability for *CCR5* to be a safe harbor for the insertion and expression of therapeutic genes has been described ^22,27^. For LSDs like MPSI, the use of the safe harbor has several advantages compared to genetic correction of the affected locus: 1) it enhances therapeutic potential, as it allows for supra-endogenous expression, 2) it circumvents design for specific mutations, 3) the coding sequences can be engineered with enhanced therapeutic properties, e.g., crossing the blood brain barrier^28^, and 4) it is versatile and easily adaptable to other LSDs.

Our approach attempts to commandeer the patient’s own hematopoietic system to express and deliver lysosomal enzymes. Because of the unique ability of this system to generate tissue macrophages that can migrate into affected tissues to deliver the enzyme^29^, an HSCT-based approach will likely be more effective that other potential enzyme depots^29^ for harder-to-treat organs like the CNS and bone. The autologous source for this approach also improves on safety, by eliminating the morbidity of graft rejection, graft-versus-host disease, and immunosuppression, and can lead to earlier intervention by obviating the need for donor matching.

We studied the self-renewal and multi-lineage differentiation capacity of the modified cells to establish the potential of genome-edited HSPCs as one-time therapy for MPSI. Our data demonstrates that this approach can modify cells with long-term repopulation potential and preserves multi-lineage differentiation capacity *in vivo* and *in vitro*. In experiments comparing engraftment potential of the YFP-and YFP+ cells, as well as in bulk transplantation experiments, cells that underwent HR-GE had approximately a 5-fold lower long-term engraftment capacity. This is not entirely surprising, as HR efficiencies are higher in cycling cells^30,31^ and therefore would be expected to be lower in stem than progenitor cells. The lower engraftment could also represent a negative effect of expression of a foreign fluorescent protein in HSCs^32^ as previously substituting a truncated form of the low-affinity nerve growth factor receptor resulted in higher engraftment frequencies than using a fluorescent protein to mark HR-GE cells^15^. As observed in allo-HSCT, this engraftment challenge might be circumvented by using larger doses of genome edited cells. This can be achieved through selection followed by *in vitro* expansion in optimized culturing conditions that could help maintain self-renewal capacity, perhaps including recently discovered small molecules such as UM171^33^ and SR1^34^.

*Ex vivo* manipulation of the HSPCs allows for a thorough examination of the genotoxicity and the magnitude of biochemical potency of the cells before delivering the engineered cell drug product to patients. Through a bioinformatics-guided strategy we identified four potential off-target sites with minimal off-target activity. Fortunately, all were reversed by using a higher fidelity nuclease^25^. The conclusion that our genome editing strategy is safe is also supported by the lack of tumorigenicity in 191 mice transplanted with 90 million edited cells.

We examined the therapeutic potential of the edited HSPCs in a new model of MPSI capable of robust human cell engraftment. Engraftment of the IDUA-HPSCs led to reconstitution of enzyme activity and decreased GAG storage in multiple organs. Notably, small changes in circulating and tissue IDUA lead to significant phenotypic improvements. This is not surprising as even a small fraction of normal IDUA activity can dramatically improve the physical manifestations of MPSI. Mean IDUA activity in fibroblasts from patients with severe MPSI is 0.18% (range 0–0.6), while 0.79% residual activity (range 0.3–1.8) results in mild disease (minimal neurological involvement and the possibility of a normal life span)^35^. In fact, healthy individuals can be found with enzymatic activity as low as 4%^36^. Our data constitutes the first study to show symptomatic correction of an LSD with human genome-edited HSPCs and provides support for the further development of this strategy for the treatment of the visceral, skeletal, and neurological manifestations in MPSI.

In sum, these pre-clinical studies provide proof-of-concept evidence of the safety and efficacy of using genome-edited human HSPCs modified to express a lysosomal enzyme to correct the biochemical, structural, and behavioral phenotype of a mouse model of MPSI, a canonical lysosomal storage disease. Moreover, this work provides specific evidence of safety and efficacy to support the optimization and development of this strategy into a clinical protocol to treat patients with MPSI and a platform approach to potentially treat other lysosomal storage disorders.

## Methods

### AAV donor plasmid construction

The CCR5 donor vectors have been constructed by PCR amplification of ∼ 500 bp left and right homology arms for the CCR5 locus from human genomic DNA. SFFV, PGK, IDUA sequences were amplified from plasmids. Primers were designed using an online assembly tool (NEBuilder, New England Biolabs, Ipswich, MA, USA) and were ordered from Integrated DNA Technologies (IDT, San Jose, CA, USA). Fragments were Gibson-assembled into a the pAAV-MCS plasmid (Agilent Technologies, Santa Clara, CA, USA).

### rAAV production

We followed a protocol that has been previously reported with slight modifications^1^. Briefly, HEK 293 cells are transfected with a dual-plasmid transfection system: a single helper plasmid (which contains the AAV rep and cap genes and specific adenovirus helper genes) and the AAV donor vector plasmid containing the ITRs. After 2 days the cells are lysed by three rounds of freeze/thaw, and cell debris is removed by centrifugation. AAV viral particles are purified by ultracentrifugation in iodixanol gradient. Vectors are formulated by dialysis and filter sterilized. Titers are performed using droplet digital PCR. Alternatively, viruses were amplified and purified by Vigene Biosciences (Rockville, MD, USA).

### Electroporation and transduction of cells

CCR5 sgRNA was purchased from TriLink BioTechnologies (San Diego, CA, USA) and was previously reported^2^. The sgRNA was chemically modified with three terminal nucleotides at both the 5′ and 3′ ends containing 2′ O-Methyl 3′ phosphorothioate and HPLC-purified. The genomic sgRNA target sequence (with PAM in bold) was: CCR5: 5’-GCAGCATAGTGAGCCCAGAA**GGG**-3’. Cas9 protein was purchased from Integrated DNA Technologies. RNP was complexed by mixing Cas9 with sgRNA at a molar ratio of 1:2.5 at room temperature. CD34^+^ HSPCs were electroporated 2 days after thawing and expansion by using the Lonza Nucleofector 4D (program DZ-100) in P3 primary cell solution as follows: 10 × 10^6^cells/ml, 300 µg/ml Cas9 protein complexed with 150 µg/ml of sgRNA, in 100 µl. Following electroporation, cells were rescued with media at 37°C after which rAAV6 was added (MOI 15,000 of 15,000 titrated to maximize modification efficiency and cell recovery). A mock-electroporated control was included in most experiments where cells underwent electroporation without Cas9 RNP.

### Quantification of putative CCR5 gRNA off-target activity by deep sequencing

Potential off-target sites in the human genome (hg19) were identified and ranked using the recently developed bioinformatics program COSMID, allowing up to three base mismatches without insertions or deletions and two base mismatches with either an inserted or deleted base (bulge). The top ranked sites were further investigated. Off-target activity at a total of 67 predicted loci was measured by deep sequencing in two biological replicates of CB-derived HSPCs. Bioinformatically predicted off-target loci were amplified by two rounds of PCR to introduce adaptor and index sequences for the Illumina MiSeq platform. All amplicons were normalized, pooled and quantified using the PerfeCTa NGS quantification kit per manufacturer’s instructions (Quantabio, Beverly, MA, USA). Samples were sequenced using a MiSeq Illmina using 2 x 250bp paired end reads. INDELs were quantified as previously described^3^.

### Measuring insertions at the CCR5 locus with ddPCR

Genomic DNA was extracted from either bulk or sorted populations using QuickExtract DNA Extraction Solution. For droplet-digital PCR (ddPCR), droplets were generated on a QX200 Droplet Generator (Bio-Rad) per manufacturer’s protocol. A HEX reference assay detecting copy number input of the *CCRL2* gene was used to quantify the chromosome 3 input. The assay designed to detect insertions at *CCR5* consisted of: F:5’-GGG AGG ATT GGG AAG ACA -3’, R:5’-AGG TGT TCA GGA GAA GGA CA-3’, and labeled probe: 5’-FAM/AGC AGG CAT /ZEN/GCT GGG GAT GCG GTG G/3IABkFQ-3’. The reference assay designed to detect the *CCRL2* genomic sequence: F:5’-CCT CCT GGC TGA GAA AAA G -3’, R:5’-GCT GTA TGA ATC CAG GTC C -3’, and labeled probe: 5’-HEX/TGT TTC CTC /ZEN/CAG GAT AAG GCA GCT GT/3IABkFQ-3’. The accuracy of this assay was established with genomic DNA from a mono-allelic colony (50% allele fraction) as template. Final concentration of primer and probes was 900 nM and 250 nM respectively. 20 µL of the PCR reaction was used for droplet generation, and 40 µL of the droplets was used in the following PCR conditions: 95° -10 min, 45 cycles of 94° -30 s, 57°C – 30 s, and 72° -2 min, finalize with 98° -10 min and 4°C until droplet analysis. Droplets were analyzed on a QX200 Droplet Reader (Bio-Rad) detecting FAM and HEX positive droplets. Control samples with non-template control, genomic DNA, and mock-treated samples, and 50% modification control were included. Data was analyzed using QuantaSoft (Bio-Rad).

### HSPC Selection and Culturing

Human CD34+ HSPCs mobilized peripheral blood purchased from AllCells (Alameda, CA, USA) and thawed per manufacturer’s instructions. CD34+ HSPCs were purified from umbilical cord blood collected donated under informed consent via the Binns Program for Cord Blood Research at Stanford University and used without freezing. In brief, mononuclear cells were isolated by density gradient centrifugation using Ficoll Paque Plus. Following two platelet washes, HSPCs were labeled and positively selected using the CD34+ Microbead Kit Ultrapure (Miltenyi Biotec, San Diego, CA, USA) per manufacturer’s protocol. Enriched cells were stained with APC anti-human CD34 (Clone 561; Biolegend, San Jose, CA, USA) and sample purity was assessed on an Accuri C6 flow cytometer (BD Biosciences, San Jose, CA, USA). Cells were cultured at 37°C, 5% CO2, and 5% O2 for 48 hours prior to gene editing. Culture media consisted of StemSpan SFEM II (Stemcell Technologies, Vancouver, Canada) supplemented with SCF (100 ng/ml), TPO (100 ng/ml), Flt3-Ligand (100 ng/ml), IL-6 (100 ng/ml), UM171 (35nM), and StemRegenin1 (0.75 mM).

### Colony-forming unit assay and clonal genotyping

Cells were single-cell sorted into 96-well plates (Corning) pre-filled with 100 µl of methylcellulose (Methocult, StemCell Technologies).

Single YFP+, YFP-, and mock-treated cells were sorted into methylcellulose media containing SCF, IL3, erythropoietin and GM-CSF, conditions that support the growth of blood progenitor cells: erythroid progenitors (burst forming unit-erythroid or BFU-E, and colony-forming unit-erythroid or CFU-E), granulocyte-macrophage progenitors (CFU-GM), and multi-potential granulocyte, erythroid, macrophage, megakaryocyte progenitor cells (CFU-GEMM).

After 14 days, colonies were counted and scored as BFU-E, CFU-M, CFU-GM and CFU-GEMM per the manual for ‘Human Colony-forming Unit (CFU) Assays Using MethoCult’ from StemCell Technologies. For DNA extraction from 96-well plates, PBS was added to wells with colonies, and the contents were mixed and transferred to a U-bottomed 96-well plate. Cells were pelleted by centrifugation at 300xg for 5 min followed by a wash with PBS. Finally, cells were resuspended in 25 µl QuickExtract DNA Extraction Solution (Epicentre, Madison, WI, USA) and transferred to PCR plates, which were incubated at 65°C for 10 min followed by 100°C for 2 min. For *CCR5*, a 3-primer PCR was set up with a forward primer outside the left homology arm (5’-CACCATGCTTGACCCAGTTT-3’), a forward primer binding the poly-adenylation signal in all inserts (5’-CGCATTGTCTGAGTAGGTGT-3’), and a reverse primer binding inside the right homology arm (5’-AGGTGTTCAGGAGAAGGACA-3’). Accupower premix was used for PCR reaction and cycled at the parameters: 95° -5 min, and 35 cycles of 95° -20 s, 72°C – 60 s. DNA fragments were detected by agarose gel electrophoresis.

### Macrophage differentiation and flow cytometry

CD34+ HSPCs were seeded at a density of 2×105 cells/mL in untreated 6-well polystyrene plates in differentiation medium (SFEM II supplemented with SCF (200 ng/ml), Il-3 (10 ng/mL), IL-6 (10 ng/mL), FLT3-L (50 ng/mL), M-CSF (10 ng/ml), penicillin/streptomycin (10 U/mL), and cultured at 37 °C 5% CO2, and 5% O2. After 48 hours, non-adherent cells were removed from plates and reseeded in new non-treated 6-well polystyrene plates at 2×105 cells/mL in differentiation medium. Adherent cells were maintained in the same plates in maintenance medium (RPMI supplemented with FBS (10% v/v), M-CSF (10 ng/ml) and penicillin/streptomycin (10 U/mL). After two weeks, adherent cells, comprising terminally differentiated macrophages, were harvested by incubation with 10 mM EDTA and gentle scraping. For phenotypic analysis we harvested 1×105 cells per condition resuspended in 100 µl staining buffer (PBS containing 2% FBS and 0.4% EDTA). Non-specific antibody binding was blocked (5% v/v TruStain FcX, BioLegend, #422302) and cells were stained with 2 µl of each fluorophore-conjugated monoclonal antibody (30 minutes, 4°C, dark). Antibodies used were hCD34-APC (BioLegend #343510), hCD14-BV510 (BioLegend #301842) and hCD11b-PE (BioLegend #101208). Propidium Iodide (1 µg/mL)) was used to detect dead cells and cells were analyzed on a BD FACSAria flow cytometer.

### Transplantation of CD34^+^ HSPCs into NSG mice

Targeted cells (sorted or bulk) were transplanted four to five days after electroporation/transduction. YFP-negative (YFP-), and YFP-positive (YFP+) cells were isolated using FACS and ∼400,000 cells were transplanted intra-femorally into sub-lethally irradiated (2.1 Gy) 6 to 8-week-old mice. Approximately 1×10^6^ cells HPSCs modified with cassettes without YFP and were transplanted in bulk. Mice were randomly assigned to each experimental group and analyzed in a blinded fashion.

### Assessment of human engraftment

16-18 weeks after transplantation, samples of peripheral blood, bone marrow, and spleen were harvested from recipient mice. Samples were treated with ammonium chloride to eliminate mature erythrocytes. Non-specific antibody binding was blocked (10% vol/vol, TruStain FcX, BioLegend), cells were stained (30 min, 4°C, dark), and analyzed by setting nucleated cell scatter gates using a BD FACSAria II flow cytometer or BD FACSCanto II analyzer (BD Biosciences). Cells were analyzed based on monoclonal anti-human HLA-ABC APC-Cy7 (W6/32, BioLegend), anti-mouse CD45.1 PE-Cy7 (A20, eBioScience, San Diego, CA, USA), CD19 APC (HIB19, BD511 Biosciences), CD33 PE (WM53, BD Biosciences), anti-mouse mTer119 PE-Cy5 (TER-119, BD Biosciences), and CD3 PerCP/Cy5.5 (HiT3A, BioLegend) antibodies, and Propidium Iodide to detect dead cells. Human engraftment was defined as HLA-ABC^+^/HCD45^+^ cells.

### IDUA activity assay

IDUA enzyme activity was measured fluoremetrically using 4-methylumbelliferyl α-L-iduronide (4MU-iduronide) (LC Scientific Inc., Canada) per established assay conditions^4^. Briefly, for IDUA the 4-methylumbelliferyl-iduronide substrate is diluted with sodium formate buffer, 0.4 M, pH 3.5, to 6.6 mM concentration. 25 µL aliquots of substrate are mixed with 25 µL of cell or tissue homogenates and adjusted to a final substrate concentration of 2.5 mM. The mixture is incubated at 37 °C for 60 min, and 200 µL glycine carbonate buffer (pH 10.4) is added to quench the reaction. 4-MU (Sigma) is used to make the standard curve. The resulting fluorescence is measured using a SpectraMax M3 plate reader with excitation at 355 nm and emission at 460 nm (Molecular devices).

### Analysis of glycosaminoglycans

Urine and tissue GAGs were measured with the modified dimethylmethylene blue assay (DMB)^5^. Tissue samples (10 - 30 mg) were incubated for 3 hours at 65 °C in papain digest solution (calcium-and magnesium-free PBS containing 1% papain suspension (Sigma), 5 mM cysteine, and 10 mM EDTA, pH 7.4) to a final concentration of 0.05 mg tissue/mL buffer. 50 µL of extract was incubated with 200 µL DBM reagent (9:1 31 µM DMB stock (in formiate buffer 55 nM): 2 M Tris base). The samples were read on a microplate reader at 520 nm.

### Histology

Histology was performed by HistoWiz Inc. (histowiz.com) using standard operating procedures and fully automated workflow. Samples were processed, embedded in paraffin, and sectioned at 4µm. Sections were then counterstained with toluidine blue or Alcian blue, dehydrated and film coverslipped using a TissueTek-Prisma and Coverslipper (Sakura). Whole slide scanning (40x) was performed on an Aperio AT2 (Leica Biosystems).

### Immunocytochemistry

MPSI fibroblasts cultures cells were fixed in 4% paraformaldehyde in phosphate-buffered saline (PBS), blocked with 3% bovine serum albumin (BSA) in PBS, and stained with rabbit anti-LAMP1 (Abcam) followed by 1:500 dilutions of Alexa 488-conjugated anti-rabbit antibody (Molecular Probes). Mounting and staining of nuclei was done Vectashield with DAPI (Vector labs). Slides were visualized by conventional epifluorescence microcopy using a cooled CCD camera (Hamamatsu) coupled to an inverted Nikon Eclipse Ti microscope. Images were acquired using NIS elements software and analyzed with ImageJ.

### Computerized Tomography

High-resolution Micro-CT scans were acquired at Stanford Center for Innovation in In-Vivo Imaging (SCI^3^) using an eXplore CT 120 scanner (TriFoil imaging). Mice were anesthetized with isoflurane (Baxter Corporation, Mississauga, ON, Canada). The scans were obtained with voxel resolution of 100 µm, an energy level of 80 keV, and 360 degrees of whole mice. Microview software (Parallax innovations) was used for isosurface rendering and measurements. Skull thickness was quantified on Midsagittal images. Femur length was determined by measuring the long axis between the two epiphysis. Zygomatic bone thickness was measures on coronal sections, perpendicular to the axis of the zygoma. Bone lengths were determined using the line measurement tool in MicroView. Femurs were measured from the base of the lateral femoral condyle to the tip of the greater trochanter.

### Spontaneous locomotor activity

All behavioral experimenters were blind to the genotype of the mice throughout testing. All tests were conducted in the light cycle. In all experiments, animals were habituated to the testing room 2 h before the tests and were handled by the experimenter for three days before all the behavioral tests. For spontaneous locomotor activity, assessment took place using the open field test in a square arena (76 × 76 cm^2^) with opaque white walls, surrounded with privacy blinds to eliminate external room cues. Mice were placed in the center of the open-field arena and allowed to freely move for 10 min while being tracked by Ethovision (Noldus Information Technology, Wageningen, the Netherlands) automated tracking system. Before each trial, the surface of the arena was cleaned with Virkon disinfectant. For analysis, the arena was divided into a central (53.5 × 53.5 cm^2^) and a peripheral zone (11.25-cm wide).

### Passive Inhibitory Avoidance

The passive inhibitory avoidance test was used to assess fear-based learning and memory. We used a dual-compartment system (GEMINI system, San Diego Instruments), where lighted and dark compartments, equipped with grid floor that can deliver electrical shocks, are separated by an automated gate. On day one, each mouse was habituated to the apparatus by placing it into the lighted compartment. After 30 s, the gate opened allowing access to the dark compartment. When the mice entered the dark compartment, the gate closed and the time to cross after the gate opened is recorded (latency time). On day 2 or training day, the mice receive a 0.5 mA shock for 2 s after a 3 s delay after crossing from the lighted to the dark compartment. On day 3, or testing day, after being placed in the lighted compartment for 5 s, the gate opened allowing access to the dark compartment. The latency to enter the dark compartment was recorded. Maximum time to cross was 10 minutes.

### Marble Burying

Repetitive behavior was tested in the marble bury test. Individual mice were introduced into cages containing 20 black glass marbles (1.5 cm diameter, four equidistant rows of five marbles each) on top of bedding 5 cm deep. After 30 min under low-light conditions, mice were removed and the number of marbles that were at least half-covered was determined.

### Mice

NOD.Cg-Prkdc^scid^IL2rg^tmlWjl^/Sz (NSG) mice were developed at The Jackson Laboratory^6^. Mice were housed in a 12-h dark/light cycle, temperature-and humidity-controlled environment with pressurized individually ventilated caging, sterile bedding, and unlimited access to sterile food and water in the animal barrier facility at Stanford University. All experiments were performed in accordance with National Institutes of Health institutional guidelines and were approved by the University Administrative Panel on Laboratory Animal Care (IACUC 25065).

### NSG-IDUA^X/X^ mice

We used CRISPR/Cas9 to knock-in the W401X mutation (UniProtKB - Q8BMG0), analogous to the W402X mutation commonly found in patients with severe MPSI, into NSG mouse embryos^7^. The guide RNA target sequence was searched using crispr.mit.edu and six shortlisted guides close to the target site were first screened by using an *in vivo* assay in NIH 3T3 cells. Two guides, one each on both sides of the target site, were selected: Guide1 (5’-TTATAGATGGAGAACAACTC-3’) cleaves 4 bases upstream and Guide3 (5’-GTTGGACAGCAATCATACAG-3’) cleaves 44 bases downstream of the target site. The guides were prepared by in vitro transcription (HiScribe™ T7 High Yield RNA Synthesis Kit, E2040S, New England Biolabs) of a dsDNA template generated by annealing two oligos (with a T7 promoter in the sense oligo) followed by a standard PCR reaction. The ssODN donor DNA contained an intended point mutation leading to a STOP codon (TGG to TAG): 5’-ggtgggagctagatattagggtaggaagccagatgctaggtatgagagagccaacagcctcagccctctgcttggcttatagATGGAGAACAA/CTCT**A**GGCAGAGGTCTCAAAGGCTGGGGCTGTGTTGGACAGCAATCATA/ CAGTGGGTGTCCTGGCCAGCACCCATCACCCTGAAGGCTCCGCAGCGGCCTGGAGTAC-3’ (lower case is intron, upper case is exon, guide cut sites marked by “/” and the mutation in bold).

Mouse Zygotes were obtained by mating NSG stud males with super-ovulated NSG females. Female NSG mice 3–4 weeks of age (JAX Laboratories, stock number 005557) were super-ovulated by intraperitoneal injection with 2.5IU pregnant mare serum gonadotropin (National Hormone & Peptide Program, NIDDK), followed 48 hours later by injection of 2.5 IU human chorionic gonadotropin (hCG, National Hormone & Peptide Program, NIDDK). The animals were sacrificed 14 hours following hCG administration and fertilized eggs were collected. CRISPR Injection mixture was prepared by dilution of the components into injection buffer (5 mM Tris, 0.1 mM EDTA, pH 7.5) to obtain the following concentrations: 10 ng/µl Cas9 mRNA (Thermo Fisher Scientific, Carlsbad, CA), 10 ng/µl IDUA1F and IDUA3F guide RNA and 10 ng/µl ssODN Donor (Integrated DNA Technologies, Coraville, IA). Zygote injections and embryo transfers were performed using standard protocols described previously^8^. A total of 38 zygotes were injected, the surviving 27 zygotes were transferred, which yielded 7 live offspring. Among these a male homozygous for the mutation was used to establish the NSG-IDUA^X/X^ colony. Mice were genotyped by-PCR based amplification followed by Sanger sequencing using the following primers: GENO F: 5’-CATGGCCCTGTTGGGTGAGTAATGA-3’, and GENO R: 5’-TGTGGTACTCCAGGCCGCTG-3’.

### Statistical analysis

All statistical test including paired and unpaired t-tests, and one-way analysis of variance (ANOVA) followed by Tukey’s multiple comparisons test was performed using GraphPad Prism version 7 for Mac OS X, GraphPad Software, La Jolla California USA. Data was reported as means when all conditions passed three normality tests (D’Agostino & Pearson, Shapiro-Wilk, and KS normality test).

## Acknowledgements

We thank Stanford’s behavioral and functional neurosciences laboratory for advice with neurobehavioral studies. We also thank Stanford’s Binns Program for Cord Blood Research for providing cells. We also would like to give thanks to the members of the Porteus laboratory for input, comments, and discussion. This work was supported by the Stanford’s Child Health Research Institute (CHRI), the National Organization of Rare Disorders (NORD), the Thrasher Research Fund, and National Institute of Neurological Disorders and Stroke (NINDS, 1K08NS102398-01).

## Author contributions

N.G.-O. conceived the project, collected data, performed experiments, carried out the analyses, and wrote the manuscript; S.G. B. performed macrophage experiments, assisted with mouse studies and figure preparation; N.M. assisted with mouse colony management, transplantation and flow cytometry; R.O.B performed CCR5 guide design and validation; S.M. obtained and purified CD34+-HSPCs from donated cord-blood and assisted with secondary transplants; R.M.Q and C.B.G. designed and generated the NSG-IDUA^X/X^ mice; C.L. and G.B. performed and analyzed off-target studies; L.A. assisted with manuscript preparation and figure design; M.P.H. directed the project, assisted with experimental design, and manuscript preparation.

## Competing interests

M.H.P. is a consultant and has equity interest in CRISPR Tx, but CRISPR Tx had no input or opinions on the subject matter described in this manuscript.

## Materials and correspondence

Materials and correspondence to be addressed to Natalia Gomez-Ospina and Matthew H. Porteus.

---

**Extended Data Fig. 1:**
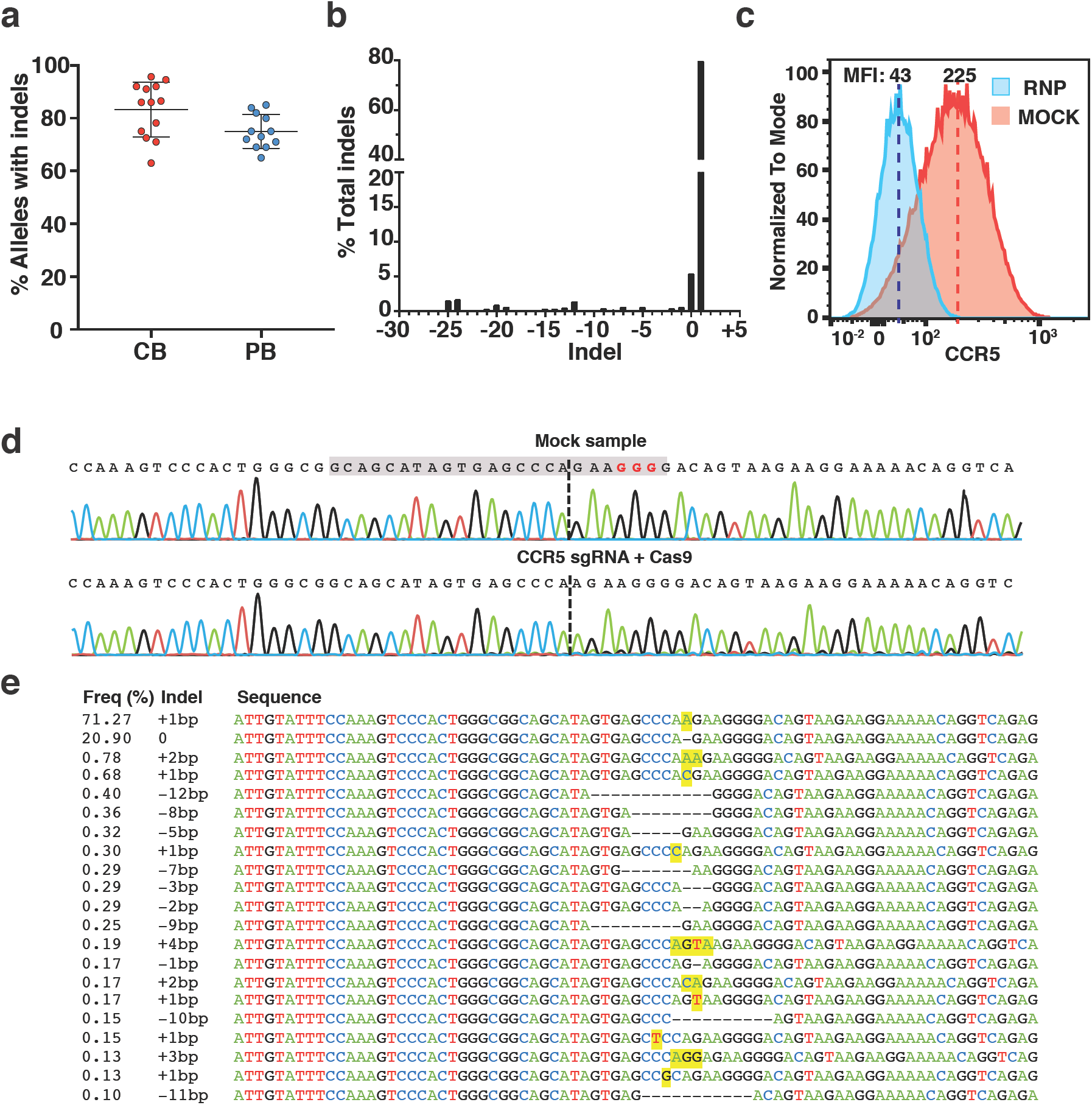
Characterization of the *CCR5* sgRNA. **a**, Indel frequency in CB and PB-derived cells by the RNP complex. **b,** Representative indel distribution from next generation sequencing reads. **c,** Histogram of CCR5 protein expression in mock-treated and RNP-treated cells showing an 80% reduction in protein expression after indel induction. **d,** Sample sequence traces around the *CCR5* sgRNA sequence (gray box, PAM in red) in mock samples and RNP-treated CB-derived HSPCs showing predominant single A insertion. **e,** Summary of indels with frequencies greater than 0.1%.

**Extended Data Fig. 2:**
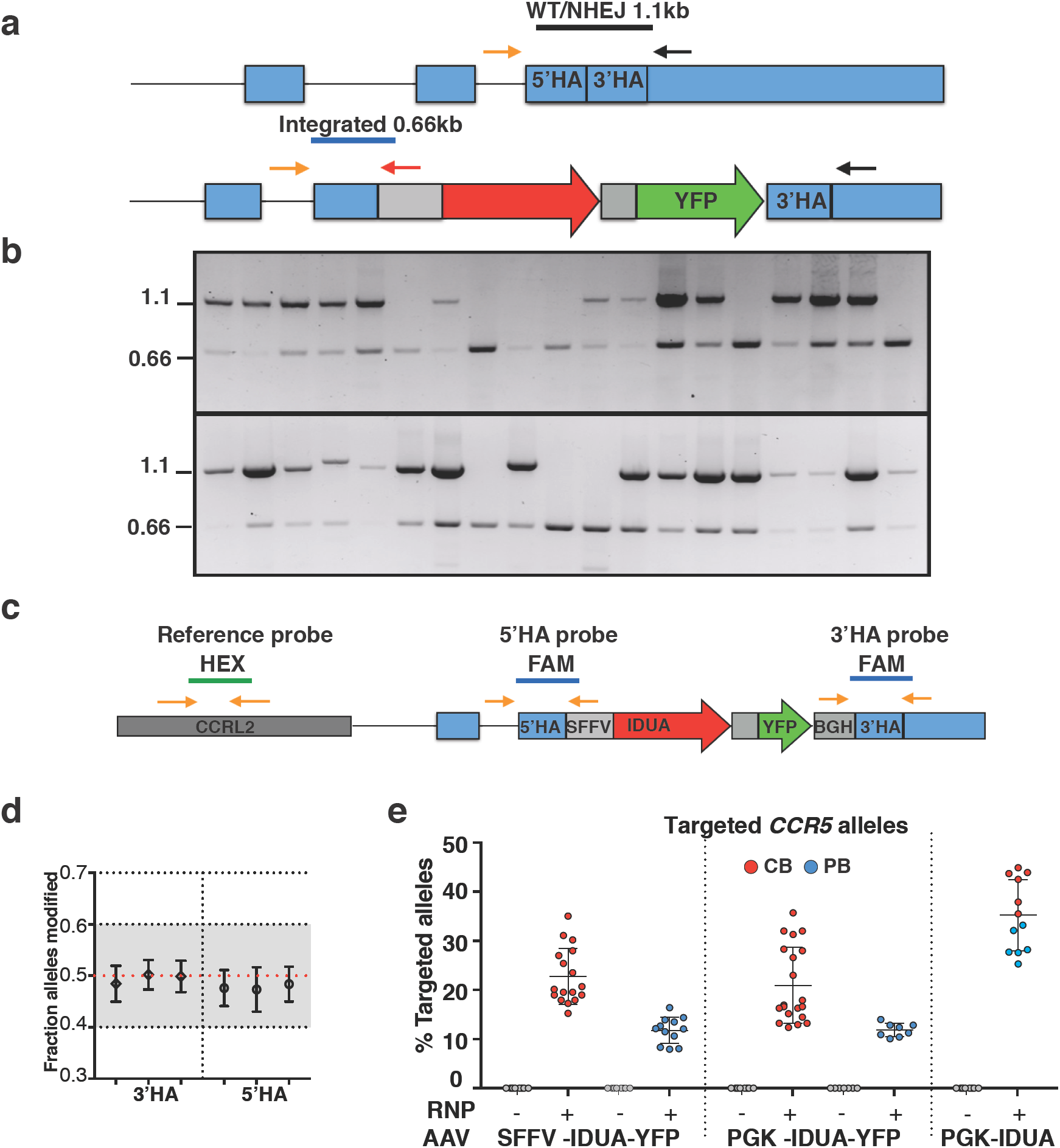
Efficiency of modification at the *CCR5* locus. **a,** Schematic showing the three primer-based genotyping scheme to distinguish mono and bi-allelic integration into the CCR5 locus on CFA-derived colonies. This strategy did not distinguish WT versus alleles with indels (NHEJ). **b,** Example agarose gels of 40 colonies genotyped in this manner. A single 1.1Kb band was interpreted as WT/NHEJ in both alleles, while a single 0.6 Kb band was read as bi-allelic integration. **c,** Schematic of probe design for ddPCR analysis. Fraction of modified alleles was obtained by using a second reference probe to the CCRL2 gene also on chromosome 3p. **d,** Originally two probes where each straddled the 5’ or 3’ homology arm were designed. The accuracy of the assays was verified and compared using genomic DNA from colonies derived from mono-allelic cells (0.5 fraction of alleles modified). Error bars indicate 95% CI. The 3’ HA probe was selected. **e,** CCR5 allele targeting frequencies in CB (red) and PB (blue)-derived IDUA expressing HSPCs as measured by ddPCR.

**Extended Data Fig. 3:**
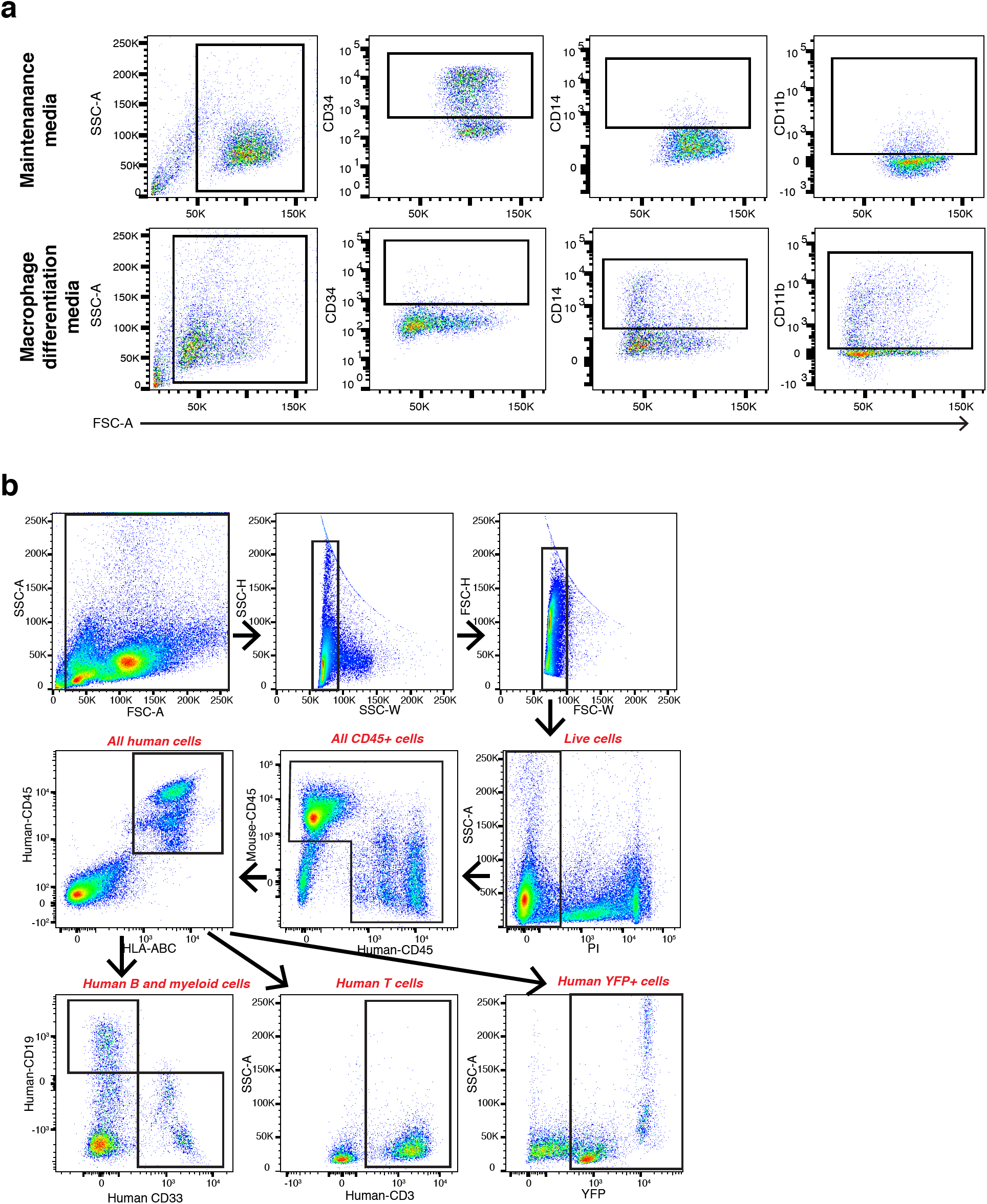
**a,** Gating scheme for quantification of human CD34+, CD14+, and CD11b+ cells in human HSPCs maintained for 2 weeks in standard CD34+ cytokine media (top panel) or media with M-CSF to induce macrophage differentiation (bottom panel). Single and live cell discrimination not shown. **b,** Gating scheme used to analyze human cell engraftment and cell lineages after transplantation. Representative plots for quantification of mouse and human hematopoietic (mCD45+ and hCD45+), all human (CD45+/HLA-ABC+), human B (CD19+), human myeloid (CD33+), human T (CD3+), and YFP+ cells.

**Extended Data Fig. 4:**
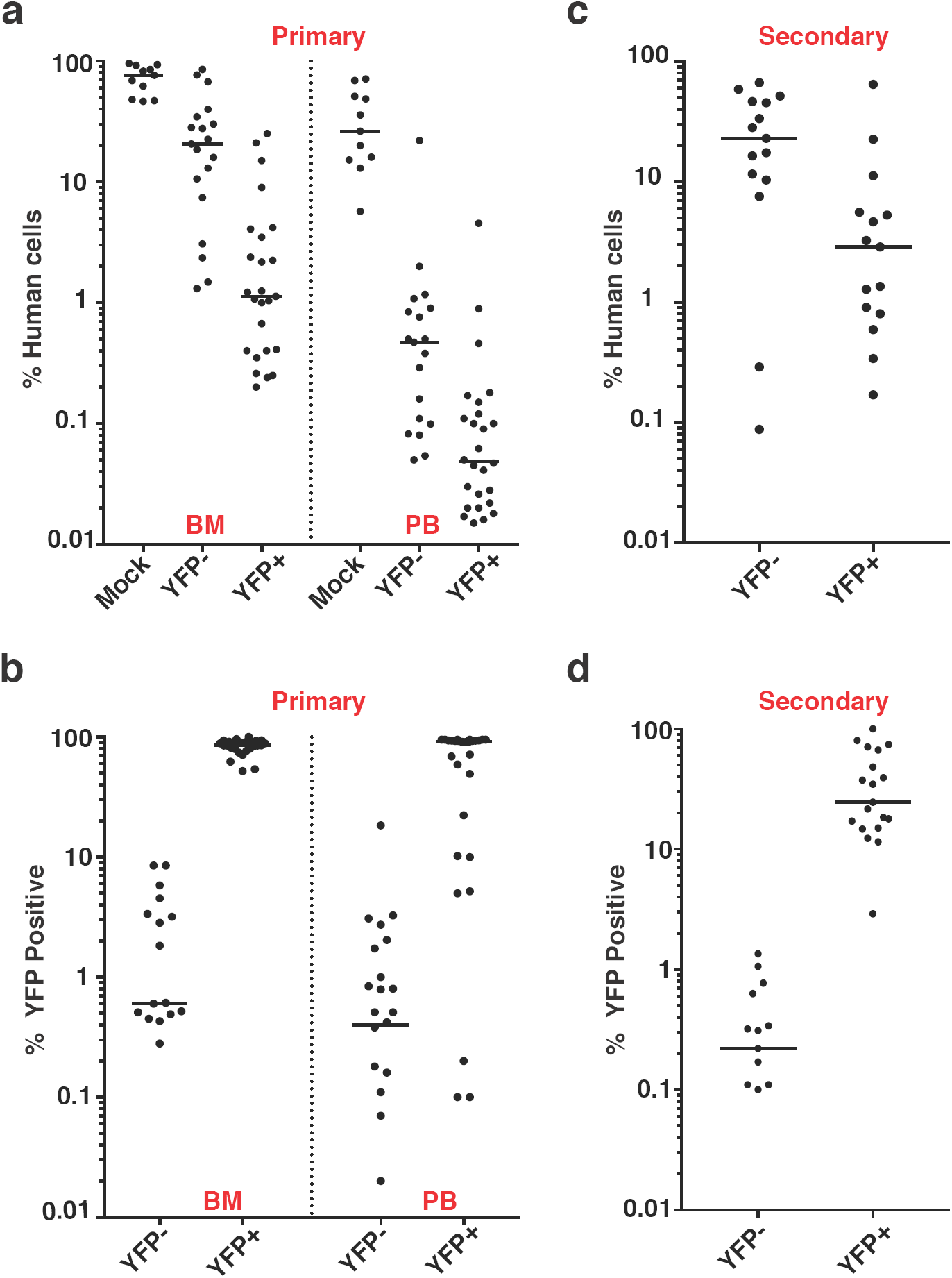
IDUA-HSPCs modified with SFFV containing cassettes are capable of long-term repopulation and multi-lineage differentiation. **a,** Percent human cell chimerism in BM and PM of mice 16-weeks post-transplant with CB-derived HPSCs. mock (n=11), YFP-(n=19), and YFP+ (n=25). Each point represents a mouse.**b,** Percent YFP+ cells in BM and PB of mice in primary transplants. **c,** Percent human cell chimerism in BM of mice in secondary transplants (32 weeks). **d,** Percent YFP+ cells in BM and PB of mice in secondary transplants. Medians shown.

**Extended Data Fig. 5:**
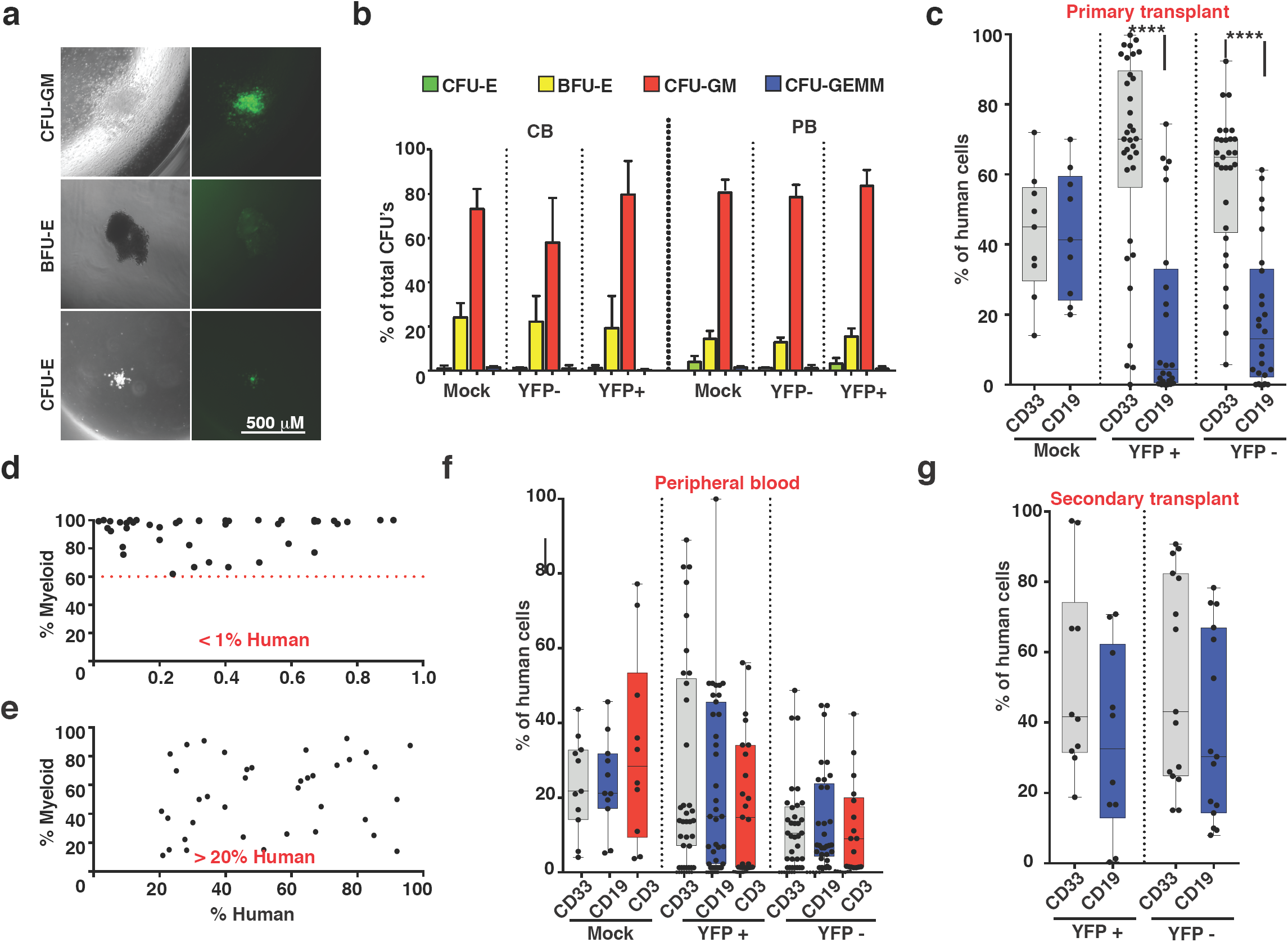
| IDUA-HSPCs maintain multi-lineage differentiation potential. **a,** Representative photos showing morphology and YFP expression in CFU-GM, BFU-E, and CFU-E colonies in CFAs. **b,** Colony formation unit frequency in mock, YFP-and YFP+ cells. **c,** Box plot with whiskers (min to max) showing percent human CD33+ (myeloid), CD19+ (B) cell in the BM of mice transplanted with mock, and FAC-sorted YFP+ and YFP-cells in primary transplant mice. Each point represents data from a single mouse. **d,** Scatter plot from mice with human cell chimerism <1% against the percent myeloid cells. **e**, Scatter plot of human cell chimerism >20% against the percent myeloid cells. **f,** Percent human CD33+, CD19+, and CD3+ (T) cells in the PB of mice transplanted with mock, and FAC-sorted YFP+ and YFP-cells. **g,** Box plot with whiskers (min to max) showing percent human CD33+ (myeloid), CD19+ (B) cell in the BM secondary transplant mice. ****: p < .0001 -in one-way ANOVA test. Post hoc comparisons were made with the Tukey’s multiple comparisons test.

**Extended Data Fig. 6:**
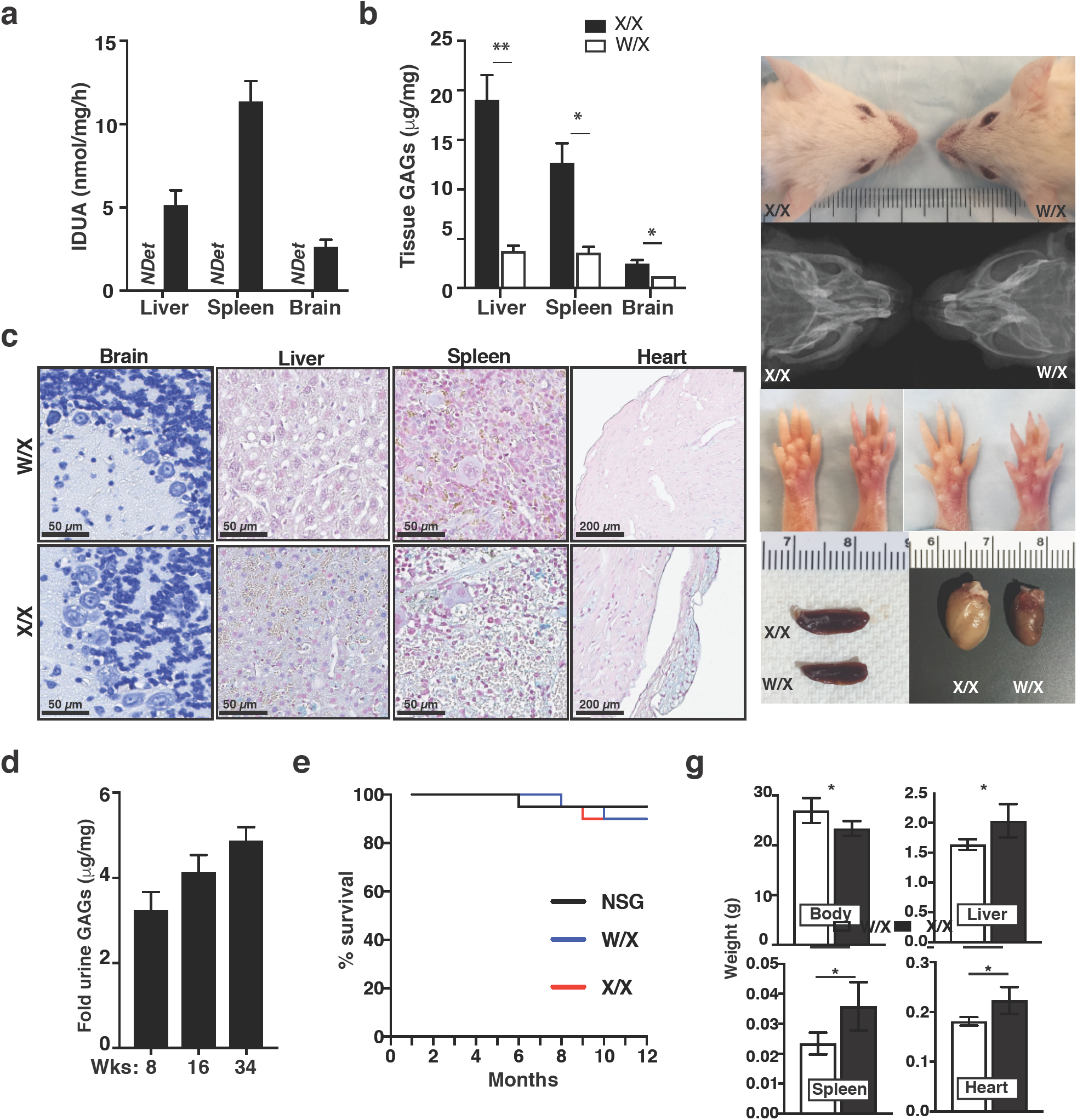
Biochemical characterization of NSG-IDUAX/X mice. **a,** Tissue IDUA enzyme activity in tissues. **b,** Tissue GAGs measure by dimethylmethylene blue reactivity. **c,** Histological sections of paraffin-embedded tissues stained with bromophenol blue (brain) or alcian blue (liver, spleen, and heart). Brain sections on X/X mice showed distended and vacuolated Purkinje cells. In liver, spleen, and heart blue deposit-laden cells can be seen throughout. **d**, Age-related progression of urinary GAGs excretion. **e,** Survival analysis comparing NSG, W/X, and X/X during one year of observation (n=10). **f,** Physical dysmorphisms and visceral enlargement. **g,** Total body, liver, spleen and heart weight in W/X and X/X mice

**Extended Data Fig. 7:**
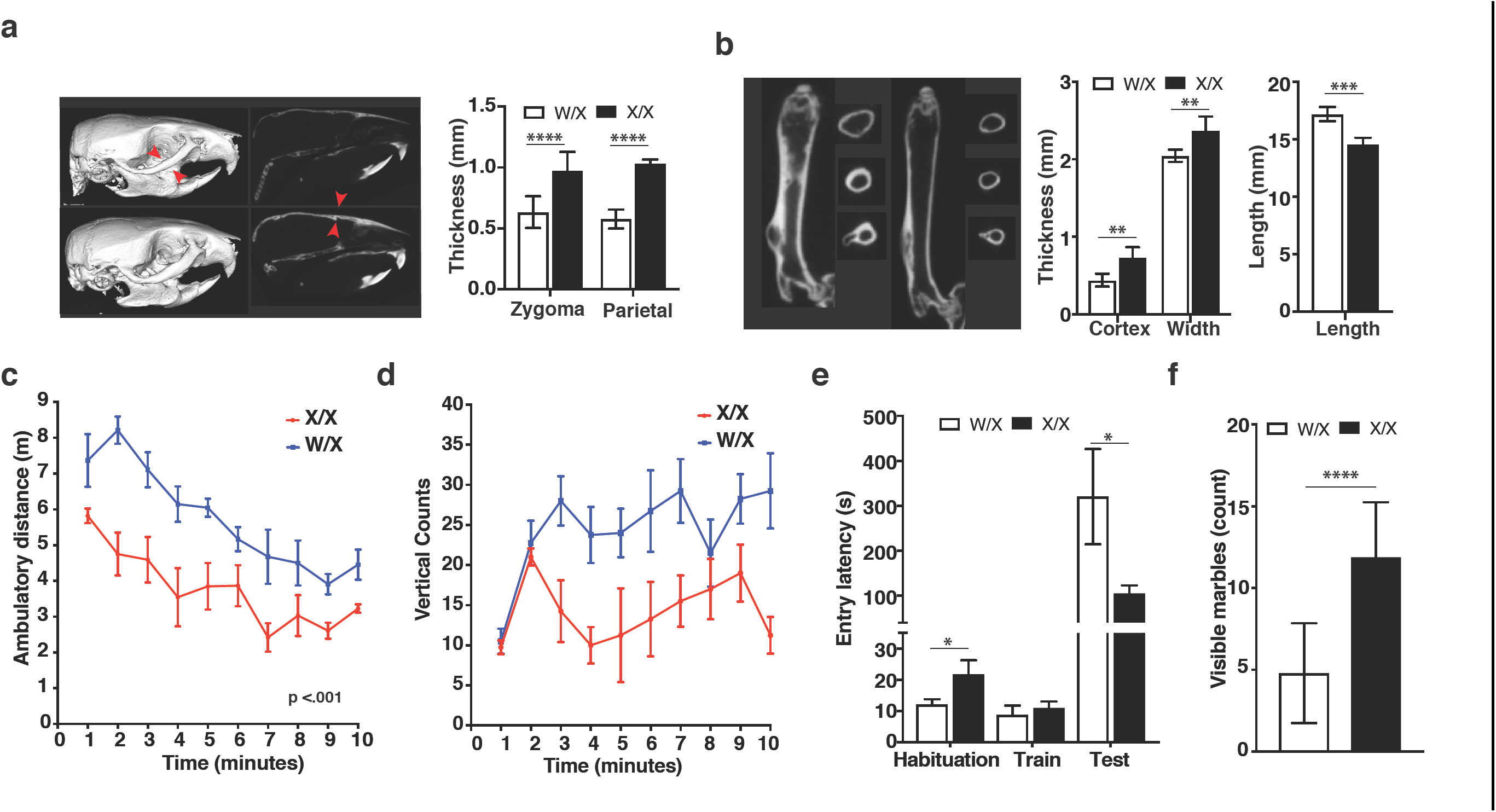
Phenotypic characterization of NSG-IDUAX/X mice. **a,** Reconstructed micro-CT images of skull and zygomatic and parietal bone thickness. **b,** CT longitudinal sections of femurs, and cortical thickness, width, and length measurements. **c,** Spontaneous locomotion in open field testing. **d,** Vertical rearing counts for W/X and X/X mice during 10-minute observation in the open field chamber. **e,** Long-term memory in passive inhibitory avoidance test. **f,** Defensive digging in the marble burying task. Total 5 female mice per genotype. Data are presented as mean ± SD for a-e, mean ± SEM for f-g. Comparisons between groups were performed using unpaired t-test. *: p < .05, **: p < .01, ***: p < .001, and ****: p < .0001. Open field testing was analyzed using within-subject modeling for the entire time course by calculating the are under the curve for each mouse and comparing between genotypes with a t-test.

**Extended Data Fig. 8:**
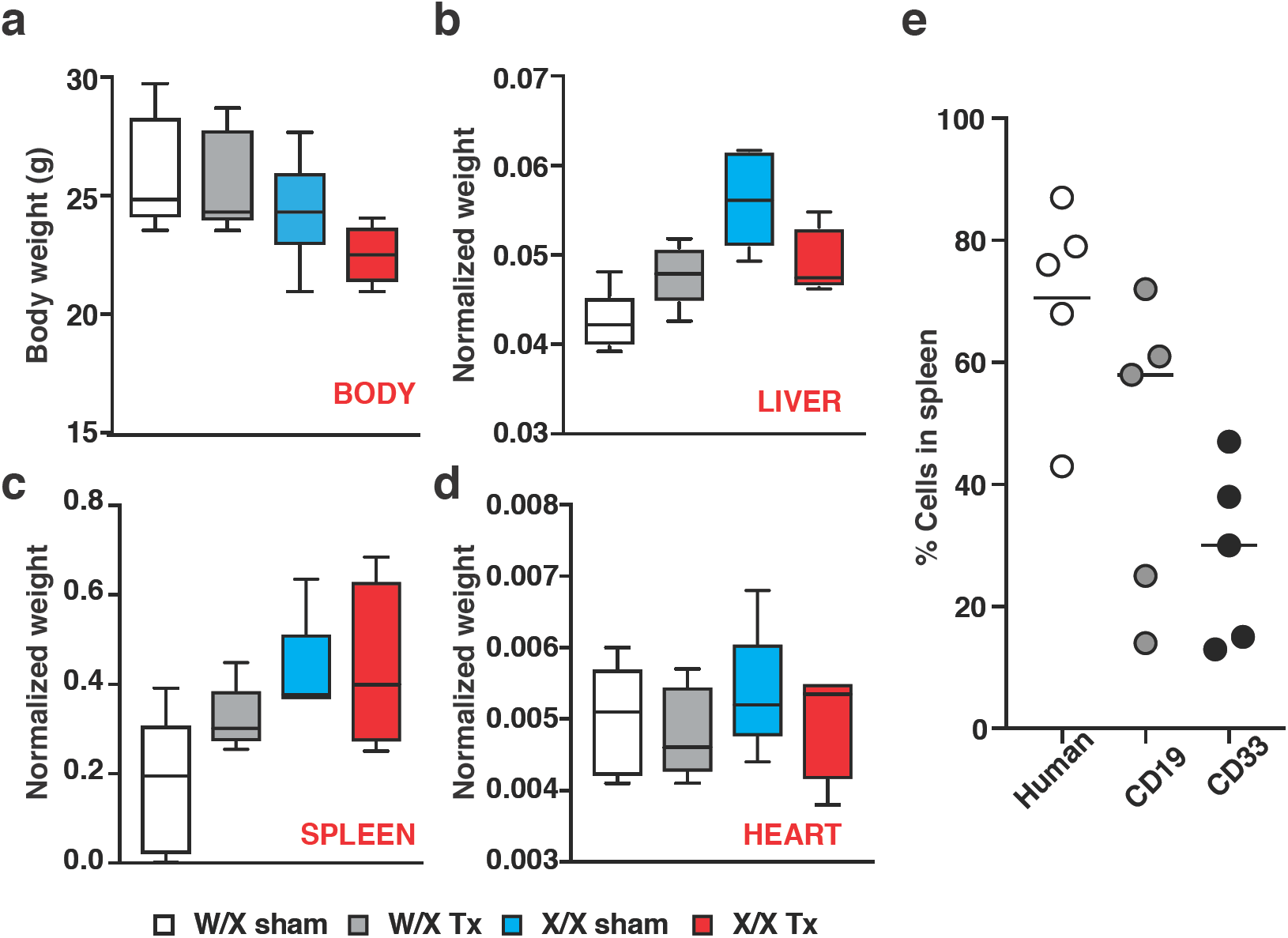
Body and organ size in transplantation experiments using bulk IDUA-HSPCs. **a**,Total body weight in heterozygous transplanted (W/X Tx-dark gray), heterozygous sham-treated (W/X sham-clear), homozygous transplanted (X/X Tx-red), and homozygous sham-treated (X/X Tx-blue) mice. **b**, Normalized liver weight. **c**, Normalized spleen weight. **d,** Normalized heart weight. **e**, Percent human, B (CD19+), and myeloid (CD33+) cells in the spleen of X/X Tx mice measured 18 weeks post-transplant.

**Extended Data Fig. 9:**
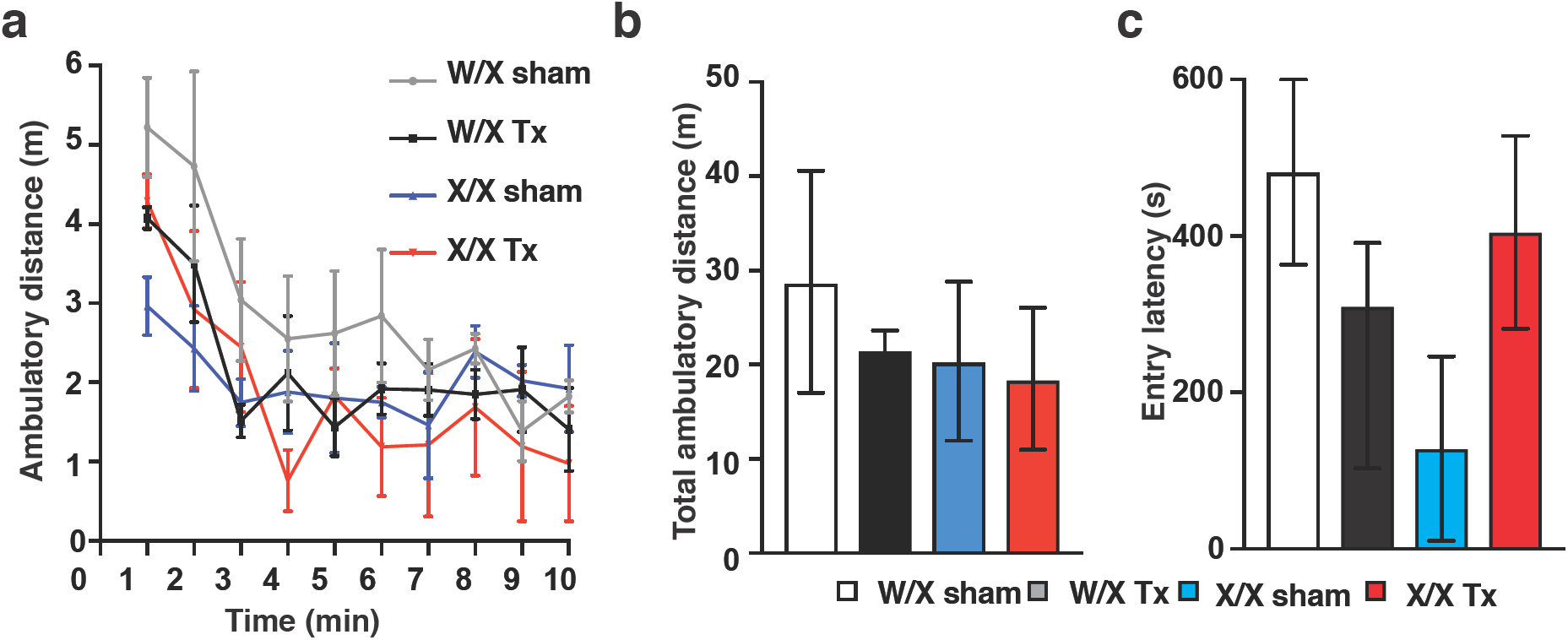
Neurobehavioral studies in mice transplanted with bulk IDUA-HSPCs. **a,** 10-minute time course of spontaneous locomotor behavior in heterozygous transplanted (W/X Tx-black), heterozygous sham-treated (W/X shamcelar or gray), homozygous transplanted (X/X Tx-red), and homozygous sham-treated (X/X Tx-blue) mice. No comparison was found to be significant. **b,** Total ambulatory distance in 10 minutes. **c,** Memory for inhibitory avoidance training (24h). No comparison was found to be significant. Comparisons between groups were performed using one-way ANOVA test and post-hoc comparisons were made with the Tukey’s multiple comparisons test. Open field testing and vertical rearings were analyzed using within-subject modeling by calculating the are under the curve for each mouse for the entire time course and comparing between groups with one-way ANOVA.

**Extended Data Fig. 10:**
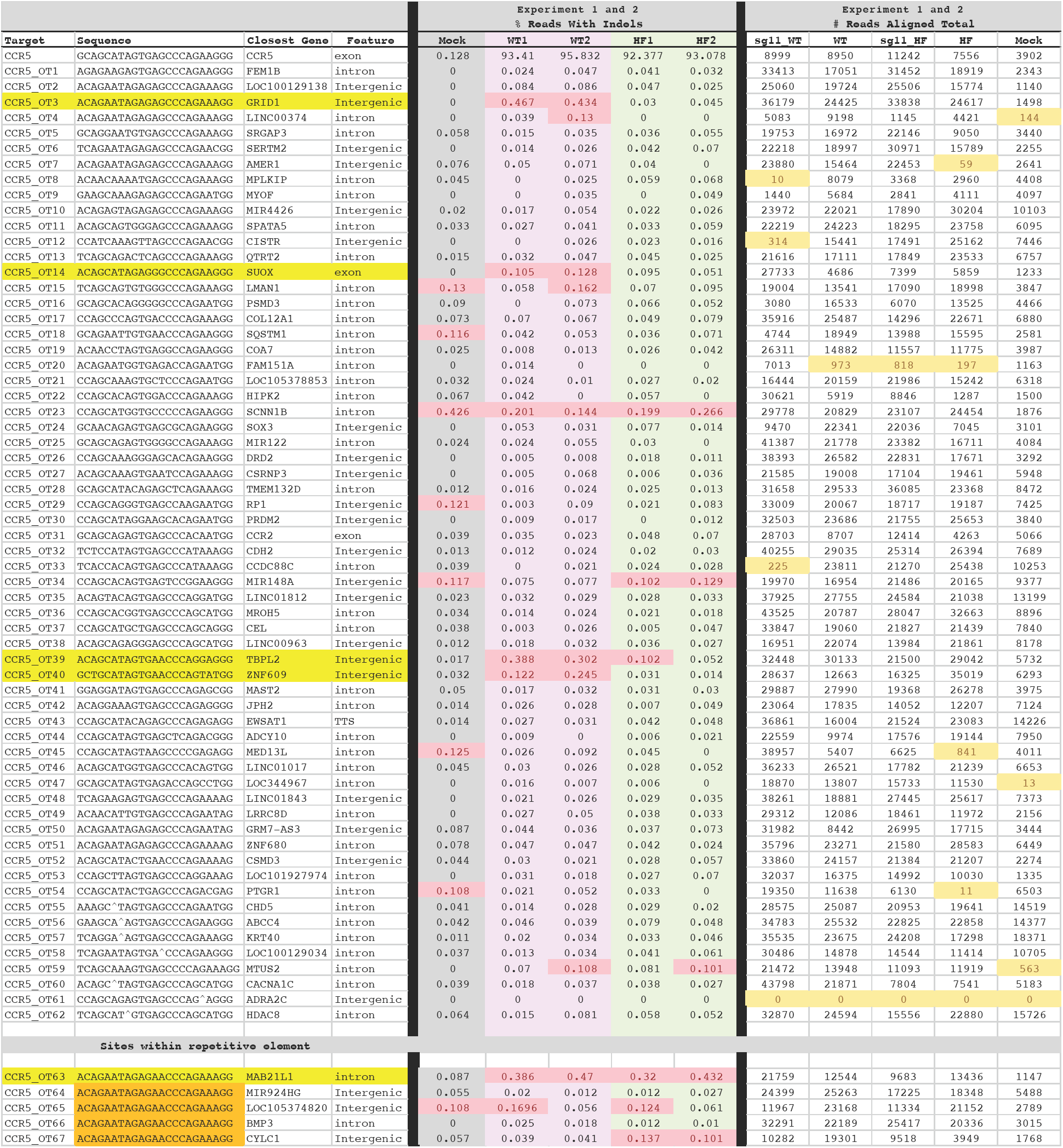
Comprehensive off-target site analysis of the CCR5 sgRNA. Table lists all 67 COSMID predicted sites, the sequence, genomic location, percent Indels in two experiments and the number of reads for each site and in each experiment. Samples with percent Indels > 0.1% are highlighted in pink. Samples with low coverage are highlighted in light yellow. CCR5 _OT63 through 67 were located within a repetitive element. Three of these had primers that should have been unique per locus but the Indel analysis showed that this was not the case. Therefore, the true off-target rated at these sites were not ascertainable but should all be < 0.5%.

